# Subtyping Abl Kinase Inhibitor Binding Modes with Machine-Learning-Enabled Super Resolution Single-Molecule Nanopore Tweezers

**DOI:** 10.64898/2026.08.27.747610

**Authors:** Ly Nguyen, Yu-Hsiang Wang, Joshua Foster, David DeCoeur, Lan Nguyen, Bryant Wu, Olgica Milenkovic, Min Chen

## Abstract

Accurate determination of kinase inhibitor binding modes could provide essential information for understanding resistance mechanisms and accelerating drug discovery. While conventional structural methods such as X-ray crystallography, cryo-EM and NMR provide high-resolution information but are low-throughput and capture largely static snapshots of dynamic protein-ligand interactions Here, we introduce a single-molecule nanopore tweezer platform that functionally subtypes ATP-competitive Abl kinase inhibitors by resolving distinct ionic current signatures of Abl-inhibitor complexes. This approach distinguishes Type I, Type IIA, and Type IIB inhibitors without structural determination. We further show how clinically relevant Abl variants (T315I and E255V) reshape inhibitor engagement and binding modes. By combining baseline probability features with wavelet-based time-frequency descriptors, ensemble machine-learning models achieved 97.5% classification accuracy across seven kinase inhibitor binding modes at sub-angstrom resolution and enabled deconvolution of mixed-inhibitor samples at nanomolar concentrations. These results establish nanopore tweezers as a label-free, super-resolution platform for profiling kinase conformational states and inhibitor binding modes, complementing structural approaches and supporting precision oncology.

## Introduction

Protein kinases are central regulators of almost all cellular signaling pathways that control proliferation, differentiation, migration, and apoptosis^1^. Abnormal kinase activity, often caused by point mutations, chromosomal translocations, or changes in expression, drives the development of many cancers such as chronic myeloid leukemia (CML)^2,3^ as well as neurodegenerative disorders including Parkinson’s and Alzheimer’s disease^4,5^. Small-molecule kinase inhibitors have emerged as one of the most successful classes of targeted therapies, achieving major clinical impact in hematological malignancies^6–8^. The discovery of imatinib^9^, followed by newer generations of drugs, demonstrated the potential of kinase inhibition and highlighted the importance of understanding inhibitor binding mechanisms. The success of targeted therapy is dramatically illustrated by the surge in approved small-molecule kinase inhibitors. As of 2026, approximately 100 protein kinase inhibitors have been approved by the U.S. Food and Drug Administration (FDA)^10^, with most of them targeting the conserved ATP-binding site. While this site is relatively accessible for inhibitor design, its high conservation across the kinome makes selectivity a major challenge and raises the risk of off-target effects^11^.

ATP-site kinase inhibitors adopt distinct binding modes within the catalytic domain and are commonly classified as Type I, Type IIA, and Type IIB based on the kinase conformation they stabilize^12^.Type I inhibitors bind the ATP pocket in the DFG-Asp-in/αC-in conformation, which is characterized by an intact regulatory spine (R-spine) - a conserved, vertical, hydrophobic, four-residue chain that bridges the N- and C-lobes and acts as a structural signature for active protein kinases^13^. In contrast, Type II inhibitors recognize a DFG-Asp-out conformation in which the R-spine is disrupted (**Fig. 1b**; **Supplementary Fig. S1**). Within this framework, Type II inhibitors are further subdivided based on their spatial occupancy: Type IIA inhibitors occupy the front cleft and gate area and extend deeply into the hydrophobic back pocket adjacent to the ATP site, whereas Type IIB inhibitors also occupy the front cleft and gate area but remain restricted from the back pocket, accessing it only partially or not at all (**Fig. 1b)**. These structurally defined binding modes strongly influence inhibitor selectivity toward resistance mutations that arise during cancer therapy, as mutations may differentially alter access to active or inactive kinase conformations. For example, the Abl T315I gatekeeper substitution blocks most Type I and II inhibitors but can be overcome by ponatinib, a type IIA inhibitor^14,15^. Consequently, determining the binding mode of a new compound at an early stage of drug development is advantageous for anticipating resistance profiles and guiding inhibitor design.

**Figure 1.**
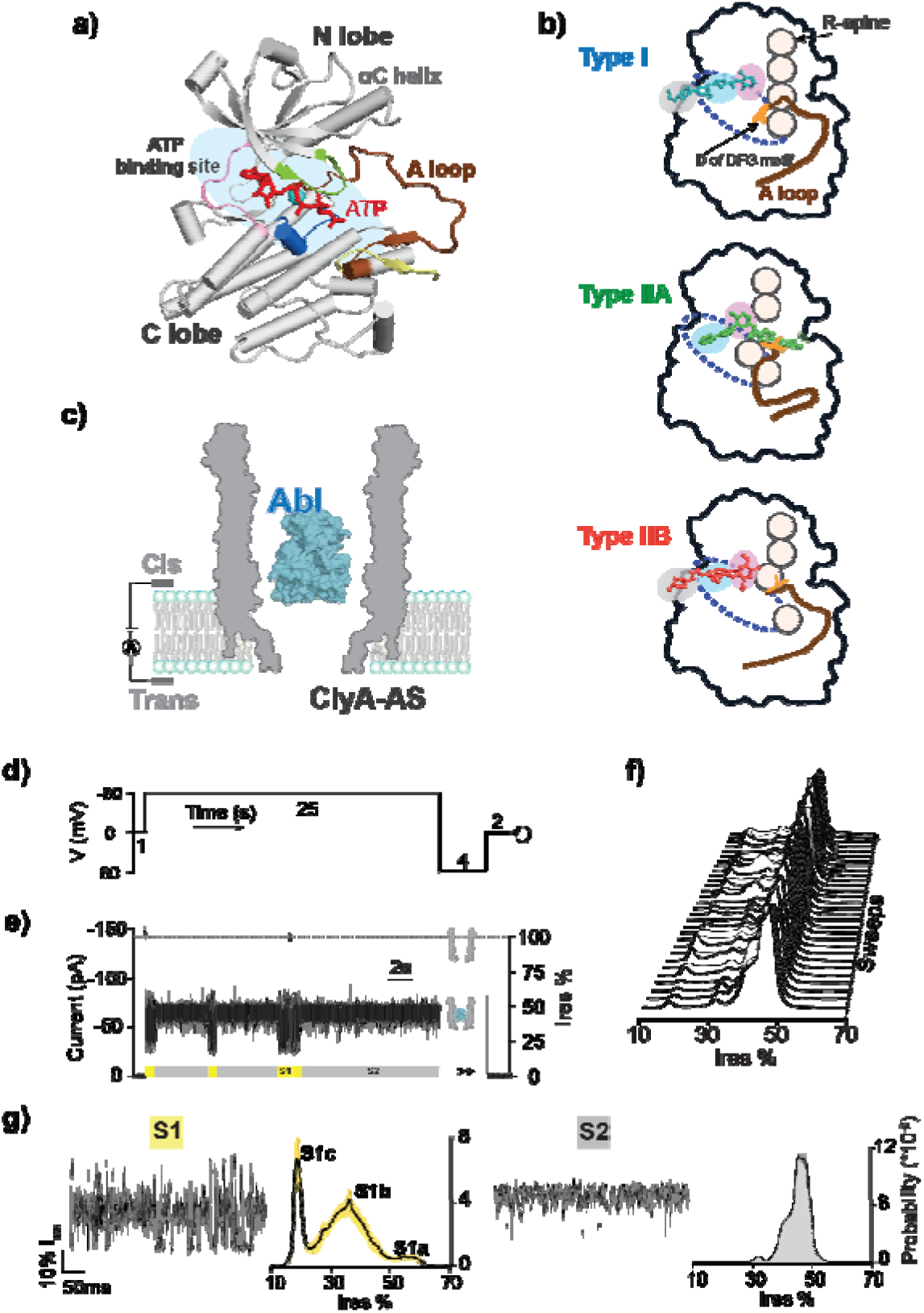
**a)** Crystal structure of human Abl catalytic domain bound with ATP-peptide conjugate (PDB: 2G1T). ATP in red and Abl substrate peptide in yellow, hinge region in pink, A-loop in brown, P-loop in green, catalytic loop in dark blue and Mg^2+^ in light blue. **b)** Schematic showing features of different ATP-site kinase inhibitor types. **c)** Schematic of ClyA nanopore tweezers for trapping Abl. **d)** A typical voltage sweep protocol; consisted of a 1s hold at 0 mV, a 25s at −80 mV, a 4s step at +80 mV to eject the trapped analyte, and a final 2s hold at 0 mV; repeated at least 60 times. **e)** A typical current trace of one voltage sweep. **f)** All point histograms of 30 randomly selected sweep traces. **g)** Zoom-in traces of S1 and S2 states and the corresponding all-point histograms of the indicated behavior averaged from 30 independent sweeps with the shade representing the standard deviation. Current recording experiments were carried out in buffer 100 mM Tris-HCl (pH 7.5),150 mM NaCl, 10 mM MgCl_2_ and 1 mM DTT.

Structural approaches, including X-ray crystallography, cryo-electron microscopy (cryo-EM) and NMR have been instrumental in defining kinase inhibitor binding modes and revealing how small molecules recognize distinct kinase conformations^7,12,16–19^. These methods have provided a foundation for structure-guided drug discovery, however, their throughput is often limited relative to the vast chemical space that need to be explored for identification of new binding modalities. As emerging resistance mutations continue to challenge targeted kinase therapies, there is a growing need for complementary approaches that can rapidly resolve binding behaviors under near-physiological solution conditions to accelerate the discovery of selective inhibitors and the development of next generation therapeutics^20^.

Nanopore analysis has emerged as a powerful single-molecule, label free platform for protein analysis, with maturing applications in protein identification^21–24^ and isoform discrimination^25,26^, ligand binding^27–29^, post-translational modification analysis^30–33^, and the study of protein-protein interactions^34–36^. In particular, nanopore tweezers have been used to monitor protein conformational dynamics in real time^37–40^. Engineered nanopores, such as cytolysin A (ClyA), and more recently YaxAB, exploit voltage-driven electroosmotic and/or electrophoretic forces to reversibly trap a single protein within a nanometer-scale cavity^41–43^ (**Fig. 1c**). The resulting ionic current fluctuations directly report protein conformational transitions and ligand-binding events with high temporal resolution and sensitivity^44^. Unlike conventional structural techniques, nanopore tweezers enable continuous, label-free monitoring of proteins in solution over timescales ranging from milliseconds to minutes^39,44^. Previously, we demonstrated that ClyA nanopore tweezers can distinguish orthosteric and allosteric kinase inhibitors based on their characteristic current signatures, highlighting their potential for mechanism-based drug classification^45^.

Here, we present a nanopore tweezer approach that subtypes ATP-site kinase inhibitors based solely on single-molecule binding signatures. Using the Abl kinase catalytic domain as a model system, we demonstrate that different types of ATP-site inhibitors produce distinct nanopore current patterns that can be classified without prior structural knowledge. We further validate the robustness of the method by applying it to clinically relevant Abl mutants and extend its application with computational algorithms for enhanced subtype discrimination and mixture analysis at nanomolar concentrations. By directly reporting on the functional dynamics of kinase-inhibitor interactions, this platform provides a rapid, label-free strategy for drug binding mode classification. More broadly, it establishes nanopore tweezers as a versatile tool for accelerating kinase drug discovery and development and supporting therapeutic development in precision oncology.

## Results and Discussion

### Nanopore tweezers resolve subtype-specific inhibitor binding signatures

Building on our previous work^39,45^, the wild-type Abl kinase domain (Abl) is driven into the ClyA-AS pore lumen under a negative applied potential (**Fig. 1c**), producing a characteristic pattern of current blockage. To efficiently sample individual Abl molecules and their ligand interactions, we implemented an episodic voltage-sweep protocol that ejects the trapped analyte every 25 seconds (**Fig. 1d**). All all-points histograms from ≥30 randomly selected sweep current traces confirmed reproducibility of kinase behavior (**Fig. 1e**). Each Abl event exhibited two distinct relative residual current states (I_res_% = 100 × I_B_/I_o_, I_B_: blocked pore current; I_o_: open pore current): S1 (14 - 62%) and S2 (30-55%), corresponding to lobe-closed and lobe-open conformations, respectively. The S1 state further resolved into three sub-states (S1a, S1b, S1c) that correspond to three ligand binding conformations, i.e. Abl:ATP binary, Abl:Abltide and Abl:ATP:Abltide ternary complexes respectively (**Fig. 1g**). As S2 was the dominant state (>95% occupancy), full-sweep histograms closely resembled that of the S2 distribution, with only minor contributions from S1 (∼5%) **(Fig. 1f & g, Supplementary Fig. S2)**.

To evaluate whether ClyA-AS nanopore tweezers can detect and discriminate the binding modes of different classes of kinase inhibitors, we tested seven inhibitors with known structural classifications (**Fig. 2**). These include five FDA-approved Abl inhibitors: the Type I inhibitor dasatinib^46^; the Type IIA inhibitors imatinib^47,48^, ponatinib^49^, and nilotinib^50,51^; and the Type IIB inhibitor bosutinib^52^. In addition, we examined vandetanib^53^, a Type I inhibitor of RET kinase, and saracatinib, a dual Src/Abl inhibitor that is not FDA-approved but has been extensively evaluated in preclinical and clinical studies^54–56^.

**Figure 2.**
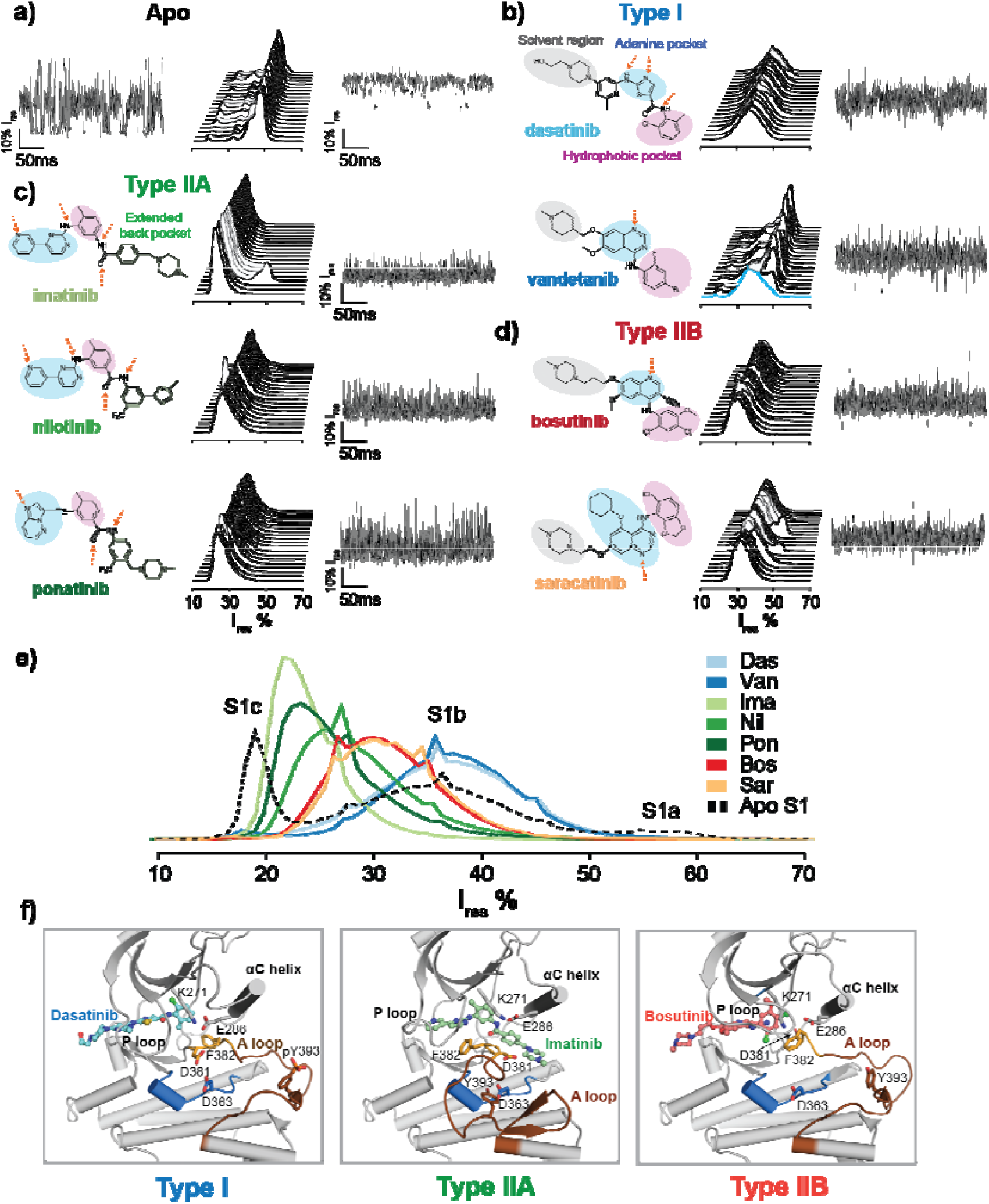
Distinct binding signals for Type I, type IIA and IIB inhibitors. **a)** All point histograms of 20 randomly selected sweep traces and representative S1 and S2 current signals of apo Abl. **b), c), d)** Chemical structures of inhibitors grouped to each type alongside their corresponding binding signals to Abl. **e)** Overlay of average histograms comparing the binding signal distributions each inhibitor. All current recordings were performed in buffer containing 100 mM Tris-HCl (pH 7.5), 150 mM NaCl, 10 mM MgCl_2_, 1 mM DTT and 150 nM inhibitors as indicated. **f)** Crystal structures of Abl kinase binding to Type I (dasatinib), Type IIA (imatinib) and Type IIB (bosutinib) inhibitors (PDB: 2GQG, 2HYY, 3UE4).

Addition of each compound resulted in a pronounced alteration of the current signal patterns **(Fig. 2b-d, Supplementary Fig. 3-9)**. Most notably, binding of all inhibitors abolished the characteristic two-state S1/S2 transitions observed for apo Abl. Instead, all inhibitor-bound events exhibited a single dominant current fluctuation pattern. This behavior is consistent with the binding mechanism of ATP-site inhibitors, whose binding is expected to stabilize the lobe-closed state. Among all inhibitors, only vandetanib showed heterogenous current patterns across different sweeps. Detailed analysis revealed that in the presence of 150 nM vandetanib, a majority of events remained in the apo-like state **(Supplementary Fig. S4)**, whereas all other inhibitors produced ∼100% of a new current signal distinct from that of apo Abl. These results indicate that vandetanib binds Abl weaker than the other inhibitors and does not fully saturate Abl at 150 nM.

Based on their current signatures, inhibitor-induced signals were grouped into three distinct categories. In the first category dasatinib, a known Type I Abl inhibitor, produced current traces with a Gaussian-like distributions centered at I_res_ ≈ 36%, closely resembling the S1b substate. Interestingly, vandetanib exhibited a highly similar current profile, with an I_res_ center around 37%. Vandetanib has been reported to adopt a Type I binding mode for RET kinase^12,53^ but has not specifically been studied in the context of Abl engagement. The observed current signature closely matched that of dasatinib, suggesting that vandetanib also engages Abl in a Type I-like binding mode. In the second group, Type IIA inhibitors, including imatinib, nilotinib, and ponatinib, generated asymmetric current distributions with substantially lower I_res_ centers (22%, 25%, and 25%, respectively), resembling an S1c-like state. Finally, the Type IIB inhibitor bosutinib displayed an intermediate signal profile, with I_res_ centered around 30%, exhibiting mixed S1b/S1c-like features and non-Gaussian statistics. Saracatinib displayed nearly identical signal characteristics (I_res_ ∼ 30%, mixed S1b/S1c-like features), suggesting that it functionally resembles a Type IIB inhibitor.

Interestingly, the signal profiles of Type I and Type IIB inhibitors share more distribution features with each other than with IIA, differing primarily by a shift from a dominant S1b state (I_res_ ∼ 36%) to mixed S1b/S1c-like features (I_res_ ∼ 30%). Although Type I inhibitors bind the active DFG-in conformation whereas Type IIB inhibitors bind the inactive DFG-out conformation, both maintain an open activation loop (A-loop) (**Fig. 2f**). In contrast, a Type IIA binding mode is associated with a closed A-loop (**Fig. 2f**). The Cα RMSD values for Type I versus Type IIA, Type I versus Type IIB, and Type IIA versus Type IIB are 3.75 Å, 1.36 Å and 3.95 Å, respectively. Thus, the similarity of nanopore current signatures among inhibitor-bound states aligns with their structural resemblance. This result can be explained by the fact that the A-loop is bulkier and more exposed to the nanopore lumen than the DFG motif, and therefore likely exerts a greater influence on the observed current signatures.

Overall, these signal profiles closely mirror the binding-mode distinctions defined in the structural classification schemes of Roskoski^12^. Importantly, the nanopore measurements complement this structural framework by revealing how inhibitors redistribute the conformational ensemble of Abl in solution. Rather than providing a single static structural snapshot, the nanopore platform continuously monitors transitions between current states, allowing ligand-induced changes in state occupancy, conformational exchange, and binding stability to be resolved at the single-molecule level. The agreement between nanopore-derived subtypes and crystallographic classifications demonstrates that ionic current signatures serve as sensitive proxies for kinase conformational states. Furthermore, the ability to functionally classify inhibitors such as saracatinib, without requiring a solved structure, highlights the potential of the nanopore tweezer platform as a rapid and label-free method for determining drug interaction mode and resistance analysis.

### Subtyping of Kinase Inhibitors Against Abl Resistance Mutants

Resistance associated mutations in BCR-Abl pose a major challenge to effective kinase inhibitor therapy in CML, often by altering kinase conformational propensity^57^ and inhibitor binding. Among these, the T315I and E255V substitutions are clinically prominent yet display distinct resistance profiles across approved inhibitors^49^ (**Fig. 3a**). For examples, T315I renders nearly all approved kinase inhibitors ineffective except ponatinib^15,58^, while E255V causes strong resistance to imatinib, moderate resistance to nilotinib, and largely retains sensitivity to other inhibitors^59^. Here, we investigate inhibitor binding modes and affinities against these two mutants.

**Figure 3.**
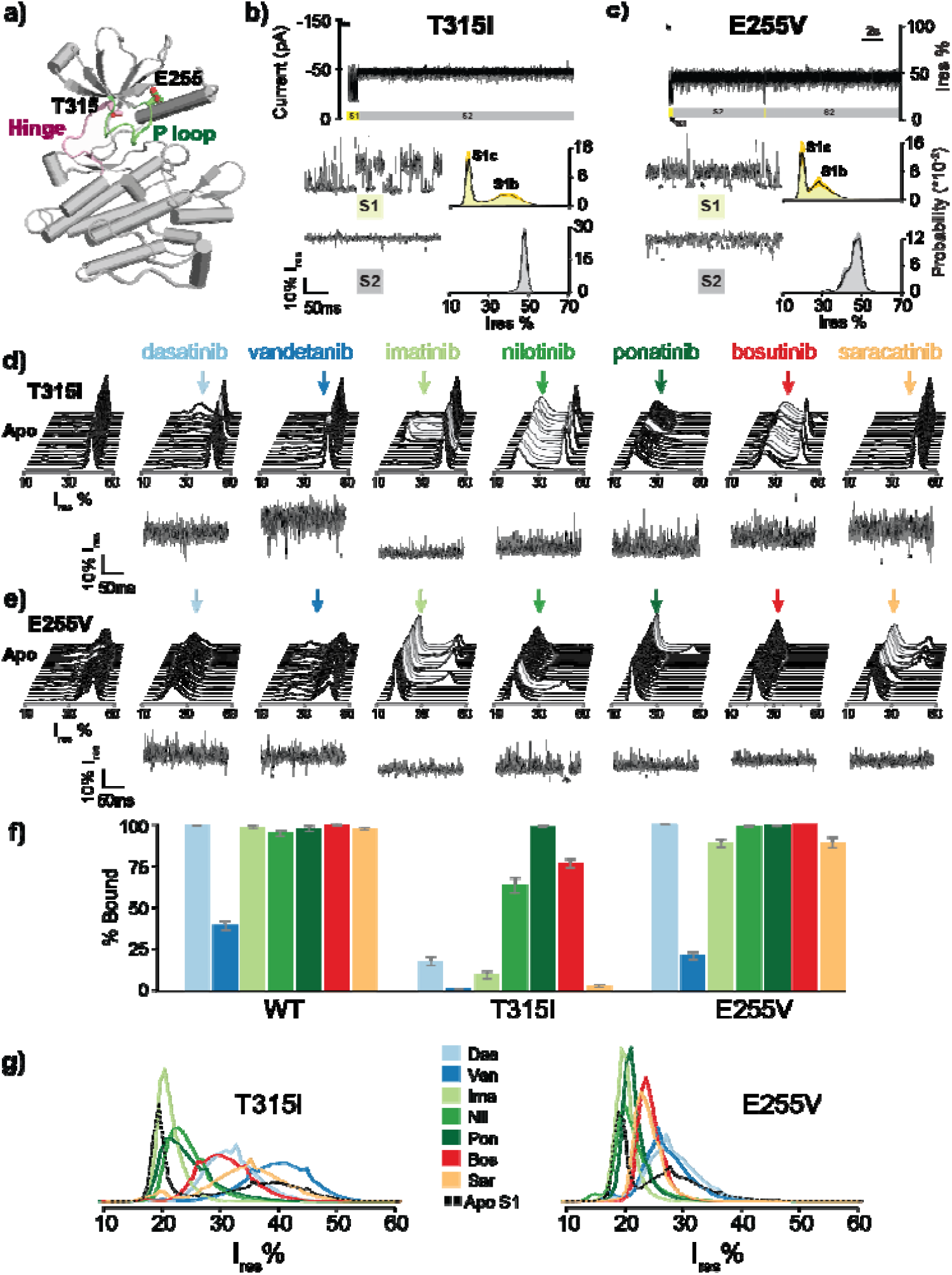
Analysis of Abl mutants trapped in ClyA nanopore tweezers. **a)** Crystal structure of Abl with highlighted T315 in the hinge region (pink) and E255 in P loop (green) (PDB: 2HYY) **b,c)** A representative trace of a single sweep for T315I (b) and E255V (c). Zoom-in traces of S1 and S2 states and the corresponding all-point histograms averaged from 30 independent sweeps of Abl mutants are also shown. **d, e)** All-point histograms of 20 randomly chosen sweeps and a representative binding signal of various inhibitors to Abl T315I and E255V mutants. Arrows indicate the histogram center of inhibitor bond events. **f)** Comparison of the percentage of drug bound events (duration weighted) for different inhibitors between Abl WT and T315I and E255V mutants, measured at 150 nM inhibitor concentration. Error bars represent the 95% bootstrap CIs (1000 iterations). **g)** Histogram overlays comparing binding signal distributions of various inhibitors to Abl T315I and E255V In contrast, the E255V mutant largely retained the canonical subtype-specific signal patterns for all inhibitors **(Fig. 3g)** For example, Type I inhibitors preferentially stabilized S1b at approximately 28% Ires, whereas Type IIA inhibitors stabilized the S1c state at approximately 20–21% Ires. Type IIB inhibitors produced intermediate profiles with mixed S1b/S1c features at approximately 24% Ires. Notably, all E255V signal distributions, including the apo signals were narrower in Ires, spanning approximately 20–28%, than those observed for wild-type and T315I Abl, which spanned approximately 20–37%. We surmise that substitution of the negatively charged glutamate with a neutral valine reduces the net negative charge of Abl, allowing the protein to be driven deeper into the ClyA nanopore under electrophoretic force, where the pore diameter is smaller. This deeper insertion likely increases steric confinement, resulting in more constrained conformations and narrower current distributions.

The apo Abl T315I (**Fig. 3b, Supplementary Fig. S10)** and E255V (**Fig. 3c, Supplementary Fig. S11)** mutants exhibited altered current profiles compared to wild-type Abl. Both mutants showed reduced duration of the S1 (lobe-closed) state, indicating global kinetic destabilization of the binding-competent conformation. In addition, T315I showed a markedly altered S2 lobe-open current profile, with a peak centered at 49% Ires and a substantially reduced width of approximately 4%. Consequently, the all point histogram of T315I sweep signals differed substantially from that of wild-type Abl (**Fig. 3d**), whereas the E255V histogram remained broadly similar to that of the wild type.

Addition of individual inhibitors triggered distinct responses. For T315I, dasatinib, vandetanib, imatinib, and saracatinib produced binding events that were distinguishable from the apo signal, but these inhibitor binding events occurred at low frequency. As a result, the sweep histograms largely resembled the apo sweeps, with rare binding events appearing sparsely in individual sweeps, as indicated by arrows. In contrast, ponatinib induced a complete signal shift, with no detectable apo-like signal. Nilotinib and bosutinib showed intermediate behavior: some sweeps displayed a clear inhibitor-induced shift, whereas others resembled the apo signal. Most inhibitors exhibited binding behavior consistent with their known clinical potencies (**Fig. 3g**). Quantification of inhibitor-bound events showed that dasatinib, which has an IC50 >1000 nM, displayed a marked reduction in binding frequency when compared to T315I, with bound events accounting for only 17.9% of the total. Imatinib, with an IC50 >6400 nM, and saracatinib showed near-complete loss of binding propensity, with bound-event fractions of 9.6% and 3.1%, respectively. In contrast, ponatinib shifted nearly the entire signal distribution to the bound state, with a bound-event fraction of 98.8%, consistent with its nanomolar potency and unique clinical efficacy against T315I (IC50 approximately 11 nM). Nilotinib and bosutinib retained moderate binding frequencies of 63.6% and 76.7%, respectively, despite clinical resistance to these inhibitors (IC50 >2000 nM). In addition to their reduced binding frequencies, dasatinib, imatinib, and saracatinib exhibited only transient interactions with T315I, with dwell-times of approximately 1s, whereas ponatinib remained bound to T315I throughout the period in which the kinase was trapped inside the ClyA lumen (**Supplementary Figs. S12–S18**). These results indicate that resistance arises from steric obstruction not only preventing productive structural accommodation but also hindering inhibitor access to the binding pocket.

By comparison, E255V exhibited only a minor reduction in imatinib binding and maintained sensitivity to the other Abl inhibitors tested (**Fig. 3e, Supplementary Figs. S19–S25**). Quantification of bound events showed that imatinib occupied E255V in 88.8% of events, compared with 98.6% for wild-type Abl, whereas all other inhibitors maintained near-complete binding, with bound-event fractions greater than 95% (**Fig. 3f**). Interestingly, these results do not fully mirror clinical data or cellular assays, in which E255V causes strong resistance to imatinib (IC50 approximately 6400 nM) and, to a lesser extent, nilotinib (IC50 approximately 430 nM)^60,61^. This is not surprising as approximately one-third of mutations classified as imatinib-resistant were previously shown to remain sensitive to imatinib and exhibited binding affinities similar to those of wild-type Abl.^62^ These observation indicate that kinase inhibitor resistance is complex and multifactorial. Although reduced inhibitor binding and loss of target engagement represent direct mechanisms of resistance, other factors - including target gene amplification, drug efflux, poor patient adherence, and changes in the tumor microenvironment - can also contribute substantially. Therefore, the strong resistance of E255V to imatinib, and its moderate resistance to nilotinib, may arise from mechanisms beyond direct binding affinity alone.

Notably, the current profiles of Type IIB inhibitors bound to T315I suggest that these inhibitors largely retained the same binding modes observed for wild-type Abl **(Fig. 3g)**. However, we also observed that the T315I mutation induced inhibitor-type switching relative to the binding modes detected with wild-type Abl. Dasatinib, which stabilizes an S1b-like Type I conformation in wild-type Abl (Ires approximately 36%), shifted to an S1b–S1c stabilization pattern in T315I (Ires approximately 31%), consistent with a Type IIB-like binding mode. Conversely, saracatinib, which exhibited an S1b–S1c stabilization pattern in wild-type Abl (Ires approximately 31%), shifted toward a more S1b-centered distribution in T315I (Ires approximately 36%), indicative of Type I-like behavior. These observations suggest that dasatinib and saracatinib can bind both DFG-in Type I and DFG-out Type II conformations, as previously reported for dasatinib^46^. It is worth noting that this type of switching occurs only between similar structures, Type I and Type IIB, as discussed previously.

Together, the results demonstrate that subtyping inhibitor binding modes using the nanopore tweezer approach is robust and reproducible for the wild-type Abl kinase and across clinically relevant drug-resistant mutants. The results show that this approach can reveal not only the intrinsic binding preferences of inhibitors, but also mutation-induced conformational effects that contribute to drug resistance. Importantly, these findings highlight the potential of nanopore technology for precision medicine where classification of inhibitor binding modes against patient-specific variants could help inform therapeutic decision-making.

### Computational algorithms improve classification accuracy

While histogram-based analysis provides an intuitive means to distinguish inhibitor subtypes, its discriminative power diminishes when compounds share similar scaffolds or when mixtures are analyzed. To overcome these limitations and enable high-throughput screening, we integrated advanced signal processing and machine learning into the nanopore signal analysis pipeline. This workflow converts raw ion current traces into quantitative molecular fingerprints through sequential preprocessing, dual-mode feature extraction, and ensemble classification (**Fig. 4a**).

**Figure 4.**
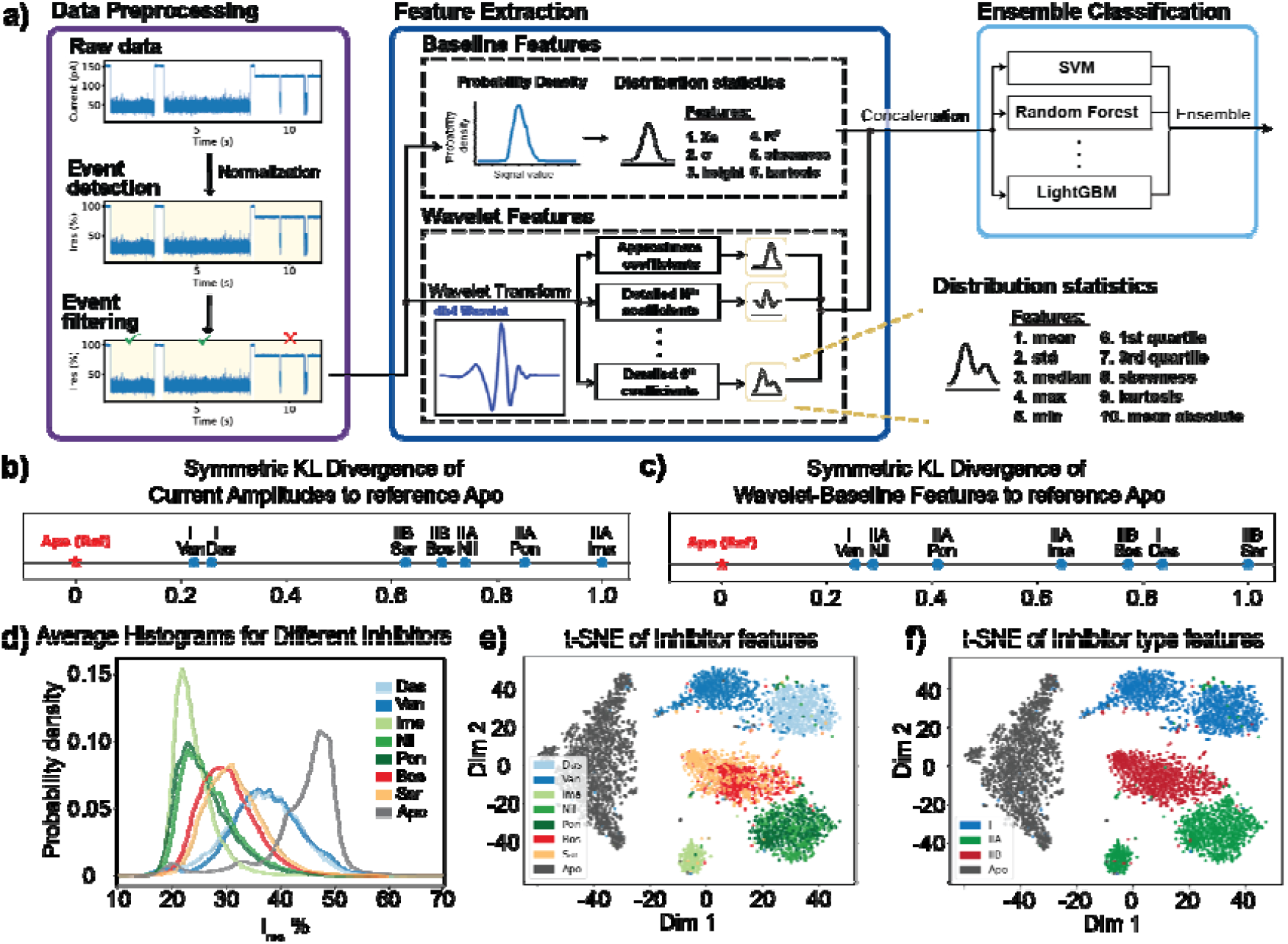
**a)** Overall workflow of the classification framework. **b)** Symmetric KL divergence of current amplitudes to reference apo. **c)** Symmetric KL divergence of wavelet-baseline features to reference apo. **d)** Probability density function of each inhibitor. **e)** t-SNE visualization of inhibitors. **f)** t-SNE visualization of binding types

To this end, raw nanopore current traces were normalized to relative residual current (I_res_) and segmented into discrete translocation events using an I_res_ = 85% threshold-based escape peak detection algorithm. To ensure data quality, events shorter than one second or with amplitudes outside the range 0–70% were removed, and gating artifacts, identified as prolonged low-amplitude signals representing partial pore blockage, were excluded.

To capture both structural and kinetic information, we implemented a dual-mode feature extraction strategy combining probability-based baseline features and time-frequency wavelet features. Baseline features were derived from the event probability density functions (PDFs), including six statistical descriptors: center of mass, standard deviation, peak height, coefficient of determination (R^2^) from Gaussian fits, skewness, and kurtosis (**Fig. 4b**). These parameters quantify signal symmetry and concentration of amplitudes, correlating with conformational states stabilized by ligand binding.

The similarity between each inhibitor and the apo reference state was quantified by the symmetric Kullback–Leibler (KL) divergence which captures differences between probability distributions (**Fig. 4c**). The symmetric KL divergence is defined as

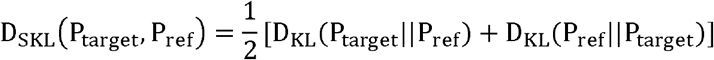

where 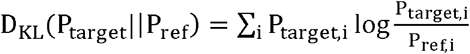 stands for the standard Kullback-Leibler divergence.

This comparison captures conformational differences between the bound and apo states. As shown in **Fig. 4b-c**, the empirical pdfs and their KL divergences provide meaningful discrimination between inhibitor types; however, compounds such as dasatinib and vandetanib remain difficult to separate based solely on amplitude distributions, prompting integration of temporal and frequency features.

Complementarily, we applied the discrete wavelet transform (Daubechies-4 basis, with four decomposition levels) to extract joint time–frequency features that capture current fluctuations associated with dynamic binding and release events. From each decomposition level, ten statistical parameters, including mean, variance, quartiles, skewness, and kurtosis, were extracted and concatenated with baseline features, producing a 56-dimensional feature space that jointly encodes distributional and time-frequency signal information.

Dimensionality reduction using the t-distributed stochastic neighbor embedding (t-SNE; perplexity = 30, learning rate = 200) revealed clear clustering of inhibitors by inhibitor feature (**Fig. 4e**) and by binding type (**Fig. 4f**). Although closely related scaffolds such as bosutinib and saracatinib or nilotinib and ponatinib showed partial overlap, distinct spatial separation among Type I, IIA, and IIB clusters confirmed that nanopore current signatures preserve subtype-specific conformational information. These analyses validate that single-molecule current dynamics contain sufficient discriminatory features to resolve fine differences between inhibitor binding modes.

To evaluate the predictive power of the extracted features, we trained a panel of supervised machine learning models and ensemble classifiers using the 56-dimensional feature set. Across all models, classification accuracy exceeded 90%, with tree-based and boosting algorithms consistently outperforming linear classifiers (**Table 1**). The stacking ensemble achieved the best overall performance, yielding 97.5% accuracy with balanced precision, recall, and F1-score of 0.975. These results demonstrate that integrating statistical and time-frequency features with machine learning enables robust, automated classification of kinase inhibitor binding modes directly from nanopore current recordings.

**Table 1.** Performance of individual classifiers and ensemble methods on the test set

| Method | Accuracy | Precision | Recall | F1-Score |
| --- | --- | --- | --- | --- |
| Individual Classifiers |  |  |  |  |
| Logistic Regression | 0.945 | 0.946 | 0.945 | 0.945 |
| SVM | 0.948 | 0.948 | 0.948 | 0.948 |
| Random Forest | 0.970 | 0.970 | 0.970 | 0.970 |
| XGBosst | 0.969 | 0.969 | 0.969 | 0.969 |
| LightGBM | 0.971 | 0.971 | 0.971 | 0.971 |
| k-NN | 0.903 | 0.905 | 0.903 | 0.902 |
| Naïve Bayes | 0.831 | 0.860 | 0.831 | 0.815 |
| MLP | 0.950 | 0.950 | 0.950 | 0.949 |
| QDA | 0.934 | 0.935 | 0.934 | 0.934 |
| Ensemble Method |  |  |  |  |
| Hard Voting | 0.970 | 0.970 | 0.970 | 0.970 |
| Soft Voting | 0.971 | 0.972 | 0.971 | 0.971 |
| Stacking | 0.975 | 0.975 | 0.975 | 0.975 |

### Classification of Mixed Inhibitor Samples

To challenge the proposed classifier approaches and evaluate their robustness and accuracy, particularly in their ability to distinguish inhibitors with similar binding characteristics in complex samples, we conducted three mixture experiments in which we used multiple inhibitors simultaneously. Due to the highly similar signal characteristics of bosutinib and saracatinib, these mixtures also tested whether scalable computational analysis could enhance signal discrimination, reduce misclassification in heterogeneous datasets, and differentiate closely related scaffolds even when binding affinity does not align with clinical potency. As shown in **Fig. 5a**, Mixture A contains all seven inhibitors (6 nM each), whereas Mixture B1 and B2 each contain six inhibitors, excluding imatinib and bosutinib, respectively. Each mixture yielded approximately 400-500 individual events encompassing both apo and inhibitor-bound segments **(Supplementary Fig. 27-29; Table S1)**. Because these composite signals often lacked clear escape peaks separating molecular states, we implemented a rolling-average low-pass filter and amplitude thresholding to divide each event into discrete segments. Each segment was then treated as an independent sample for classification **(Fig. 5b)**.

**Figure 5.**
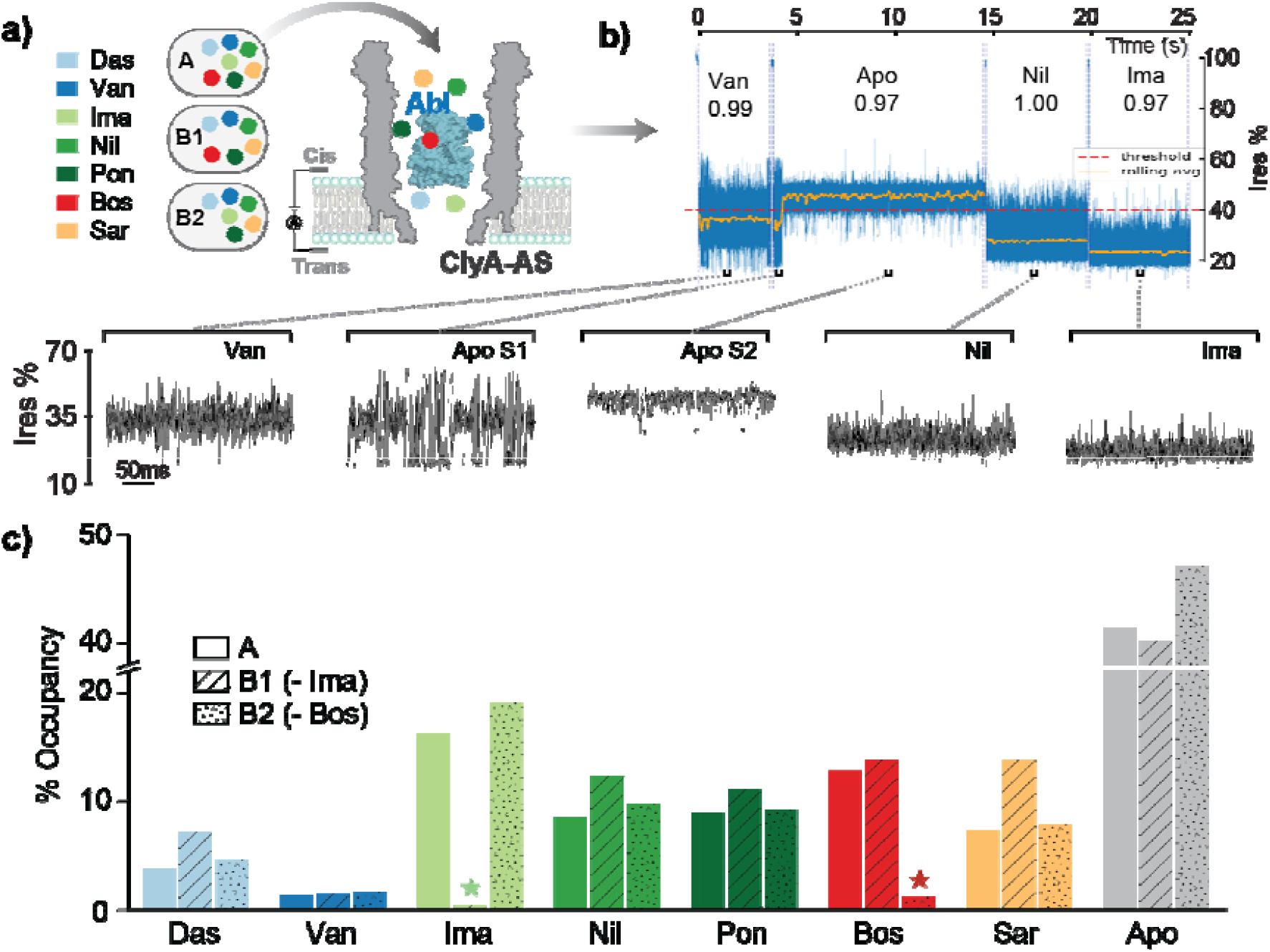
**a)** Overall workflow of the mixture experiments. **b)** A representative sweep from collected data with the predicted results and confident scores. In the mixture data, different states may not be separated by escape currents. Therefore, a rolling average with a threshold is applied to divide each single event into multiple segments, each of which is used as a unit for classification. **c)** The prediction results for the mixture data show that the classifier rarely predicts non-existent inhibitors for each mixture, demonstrating the model’s accuracy

The stacking ensemble successfully classified all inhibitors present in the mixtures, accurately identifying the absence of imatinib and bosutinib in mixtures B1 and B2, respectively, with negligible false positives (**Fig. 5c; Table S1**). Initial analyses revealed minor misassignments of apo transition events as vandetanib-binding events, attributed to partial overlap between apo S1-S2 transitions and Type I binding signatures. Incorporation of composite S1-S2 parameters and refined amplitude thresholds effectively mitigated these misclassifications, resulting in improved subtype resolution and overall accuracy across all mixtures.

By integrating probabilistic modeling, time-frequency analysis, and multivariate learning, the nanopore computational pipeline elevates molecular discrimination far beyond traditional histogram-based approaches. This framework reliably separates structurally similar inhibitors, scales for automated high-throughput analysis, and extends classical static kinase classification schemes, such as those of Roskoski and Modi & Dunbrack^12,16^, into the dynamic domain. Whereas conventional taxonomies rely on crystallographic features such as DFG-in/out or αC- in/out positions, nanopore recordings capture transient intermediates and real-time binding-state transitions, providing a functional complement to structure-based inhibitor subtyping.

This capability is particularly striking given the near-identical backbone conformations of the examined inhibitor-bound kinase structures. The Cα RMSD values between imatinib-bound Abl (PDB: 1IEP) and nilotinib-bound Abl (PDB: 3CS9), and between imatinib-bound and ponatinib-bound Abl (PDB: 3OXZ), are approximately 1.23 Å and 0.90 Å, respectively, indicating almost identical overall protein backbone architectures. Despite these sub-angstrom structural differences, ClyA nanopore tweezers, when coupled with wavelet-based feature augmentation and tailored classification methods, can robustly discriminate the three inhibitor-bound states. This demonstrates an effective super-resolution capability at the atomic level, in which subtle differences in side-chain dynamics that are difficult to distinguish by global structural comparison are amplified through nanopore readouts and learned molecular fingerprints.

Although the method has current limitations, including the observation that inhibitor binding affinities measured under the present experimental conditions do not always correlate with known clinical potencies, these discrepancies create important opportunities for future studies examining how nanopore-resolved dynamics relate to cellular activity and in vivo efficacy. Within the scope of this study, however, the primary objective is kinase inhibitor subtyping, and the results clearly demonstrate that nanopore tweezers combined with time-frequency feature extraction and learning provide a powerful and effective strategy for this purpose.

## Conclusions

Here, we demonstrate that a machine-learning-enhanced ClyA nanopore tweezer platform provides a rapid, label-free, single-molecule approach for classifying Abl kinase inhibitors by resolving distinct ionic current fingerprints associated with different inhibitor binding modes at sub-angstrom resolution. Unlike conventional biochemical and structural methods, which often report population-averaged measurements or static conformational snapshots, nanopore tweezers capture real-time conformational and kinetic equilibria with millisecond temporal resolution. Combined with machine-learning and signal processing techniques, the ClyA nanopore enables discrimination among Type I, IIA, and IIB kinase inhibitors, as well as classification of compounds with highly similar chemical scaffolds, even within mixtures. Overall, the nanopore-based classification strategy described here offers a super-resolution, highly scalable and material-efficient method for interrogating kinase-inhibitor interactions, with the potential to accelerate drug discovery and support more personalized therapeutic development.

## METHODS

### Mutagenesis, Expression, and Purification of Abl Kinase

Abl kinase variants were generated by overlapping mutagenesis PCR using pET 2BT10/His10-Tev-N4posAbl (residues 229 – 512) ^39^ as the template and primers listed in Supplementary Table S2. Plasmids contain Abl variants and YopH phosphatase (residues 164 - 468; pET13S/yopH) were co-transformed into *E. coli* BL21(DE3) pLysS cells. Co-expression with YopH phosphatase prevented Abl autophosphorylation during expression. Cultures were grown in 2xYT media supplemented with 200 µg/mL ampicillin, 50 µg/mL spectinomycin, and 30 µg/mL chloramphenicol at 30 °C with shaking at 240 rpm until the OD600 reached 0.6. Protein expression was induced with 250 µM isopropyl β-D-1-thiogalactopyranoside (IPTG) and the culture continued to grow for 16h at 16°C at 240 rpm. Cells were harvested by centrifugation at 4,000 × *g* for 20 min at 4 °C and resuspended in lysis buffer containing 50 mM Tris-HCl (pH 8.0), 150 mM NaCl, 25 mM imidazole, 100 µM PMSF, and 10% glycerol. Cells were lysed by sonication (Misonix), and the lysates were clarified by centrifugation at 50,000 × *g* for 30 min at 4 °C. The clarified supernatant was applied to a Ni-NTA affinity column containing 3 mL HisPur™ Ni-NTA Resin. The column was washed with buffer containing 50 mM Tris-HCl (pH 8.0), 150 mM NaCl, 75 mM imidazole, 100 µM PMSF, and 10% glycerol, and bound proteins were eluted with the same buffer containing 250 mM imidazole. Fractions containing Abl at high purity were pooled and incubated overnight at 4 °C with TEV protease at a 1:10 molar ratio (TEV:Abl) while being dialyzed against 50 mM Tris-HCl (pH 8.0), 150 mM NaCl, 10% glycerol, and 1 mM TCEP. The sample was subsequently passed over a second Ni-NTA column to remove His-tagged TEV protease and uncleaved His-tagged Abl. The flow-through containing tag-cleaved Abl was further purified by size-exclusion chromatography using an HW55S column (Tosoh Bioscience) equilibrated in 50 mM Tris-HCl (pH 8.0), 150 mM NaCl, 10% glycerol, and 1 mM DTT. Protein purity was assessed by 12% SDS-PAGE. Purified proteins were aliquoted, flash-frozen in liquid nitrogen, and stored at −80 °C.

### Preparation of ClyA-AS Nanopores

The pET3a/ClyA-AS-5His plasmid was transformed into *E. coli* BL21(DE3) pLysS cells. Cells were grown in 2xYT media supplemented with 200 µg/mL ampicillin and 30 µg/mL chloramphenicol at 30°C, 240 rpm until OD_600_ reached 0.6. ClyA protein expression was induced with addition of 500 µM IPTG to the culture which were continuously grown for 16h at 16°C, 240 rpm. Cells were then harvested by centrifugation (4000 g, 20 min, 4 °C) and resuspended in a lysis buffer (137 mM NaCl, 2.7 mM KCl, 10 mM Na_2_HPO_4_, 1.9 mM KH_2_PO_4_, 10% glycerol, pH 7.4) supplemented with 10 mM imidazole, and 100 µM PMSF. Cells were lysed by sonication (Misonix), and lysates were clarified via centrifugation (50,000 g, 30 min, 4 °C). The supernatants were loaded onto a Ni-NTA affinity column. The column was washed with buffer containing 20 mM imidazole, and bound proteins were eluted with buffer containing 200 mM imidazole. Elution fractions were further purified by size-exclusion chromatography using an HW55S column equilibrated in 137 mM NaCl, 2.7 mM KCl, 10 mM Na_2_HPO_4_, 1.9 mM KH_2_PO_4_, and 10% glycerol (pH 7.4) to remove protein aggregates. Protein purity was assessed by SDS-PAGE. Monomeric ClyA-AS was aliquoted, flash-frozen in liquid nitrogen, and stored at −80 °C. For ionic current recording experiments, ClyA-AS monomers were assembled into oligomers by incubation with 1% DDM at 37 °C for 30 min. The resulting oligomers were stored at 4 °C until use.

### Single-Channel current recording

Single-channel recordings were performed at 22°C using a flow cell consisting of two chambers separated by a 25 µm PTFE film (Goodfellow) containing a ∼100 µm aperture. The aperture was treated with 1 µL of 10% hexadecane (v/v in pentane). Each chamber was then filled with 600 µL of recording buffer 150 mM NaCl, 100 mM Tris-HCl (pH 7.5), and 1 mM DTT. DPhPC dissolved in pentane (18 µL, 10 mg/mL) was added to both chambers, and bilayer formation was achieved by repeatedly raising and lowering the liquid level across the aperture using a pipette. ClyA-AS nanopores were added to the grounded cis chamber and a voltage protocol was applied to facilitate the insertion. Abl kinase (∼100 nM) was added to the cis chamber with the nanopore luminal entry exposed. Gap-free recordings were initially collected for several minutes to confirm baseline stability. Afterward data were acquired using an episodic voltage protocol to measure apo and inhibitor-bound states. Each sweep consisted of a 1s hold at 0 mV, a 25s at −80 mV, a 4s step at +80 mV to eject the trapped analyte, and a final 2s hold at 0 mV; this sequence was repeated at least 60 times. All inhibitors, including dasatinib, vandetanib, imatinib, nilotinib, ponatinib, bosutinib, and saracatinib, were prepared as 100 µM stock solutions in DMSO and added to the chamber exposing the wider nanopore entry at a final concentration of 150 nM.

Ionic currents were recorded using an Axopatch 200B amplifier (Molecular Devices) and digitized with a Digidata 1440A (Molecular Devices) at a sampling rate of 10 kHz after low-pass filtering at 2 kHz with a four-pole Bessel filter. Data acquisitions were performed using pClamp 10.7. Clampfit 11.1 and OriginPro 2024 were used for data analysis. The residual current (I_res_) was calculated from the blocked pore current (I_B_) and open pore current (I_O_) as I_res_ (%) = 100 × I_B_ / I_O_. The I_res_ value at the peak center in the histograms was obtained by Gaussian multi-peak fitting using OriginPro 2024.

### Inhibitor bound percentage analysis

Determining the percentage of inhibitor bound Abl sweeps was performed via a custom Python script. First, Abl capture events were identified by continuous stretches of current between ∼0.66 and ∼0.05 I_res_ lasting for at least 500ms long. After validation of event capture events, each event’s time series current data was extracted for further analysis. To determine inhibitor bound and unbound occupancy for each Abl capture event, a two-state classification approach was used. Reference I_res_ means were dictated empirically for the inhibitor bound and unbound states and a classification threshold was set at the midpoint between these two reference levels. Each Abl capture event was smoothed using a 500ms moving average filter to reduce noise. Each point of the smoothed signal was then classified via its proximity to each reference level as either bound or unbound. To extract physically meaningful dwell-times and remove short-lived fluctuations, regions of state assignment agreement 300ms or longer were retained as valid states. Regions below this state agreement were discarded, and the samples were assigned to the nearest valid state assignment. The percentage of inhibitor bound Abl was calculated as the fraction of total samples classified as bound, resulting in a time-weighted measure of inhibitor occupancy over the duration of each capture event. To report a single measure of inhibitor-bound occupancy, an aggregated inhibitor-bound proportion was calculated as the duration-weighted mean across all events of a single condition. This method resulted in the contribution of each event being proportional to its duration. The confidence intervals for each condition were determined via nonparametric bootstrap sampling (1000 iterations). In each iteration, events were sampled with replacement, and the duration-weighted inhibitor-bound occupancy was recalculated. The 95% confidence intervals were defined by the 97.5th and 2.5th percentiles from the bootstrap-calculated means distribution.

### Data Processing and Computational Analysis

#### Data Preprocessing

Nanopore current sweeps were processed using custom Python scripts. Individual events were segmented using a threshold-based escape-peak detection algorithm applied to the residual current trace. Events shorter than 1s or with residual current amplitudes outside 0–70% I_res_ were excluded from analysis.

Gating artifacts were additionally removed using an amplitude-based filtering procedure. For each event with discrete current signal *x* [*n*], a local current amplitude sequence *A* [*n*] was computed using a sliding window of width *W* = 2000 samples (0.2s at a 10kHz sampling rate). Specifically, the local amplitude was defined as

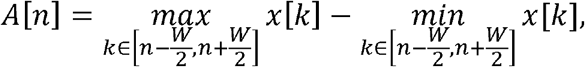

measuring the peak-to-peak variation of the current within the local window. Boundary regions of width *W /* 2 at the beginning and at the end of each event were excluded from this calculation.

Events were classified as gating artifacts if the number of samples for which the local amplitude *A* [*n*] fell below a predefined threshold exceeded a duration threshold of 5000 samples (0.5s). The amplitude threshold used was set to I_res_ = 0.20 for inhibitor-bound classes, and to I_res_ = 0.15 for the apo class, as apo events generally display lower current fluctuations. Gating events were excluded from all subsequent analysis.

#### Feature Extraction

To capture both stationary information and transient dynamics in nanopore current signals, each retained event is represented using a combined *baseline* and *wavelet feature set*. Baseline features extract distributional information from the probability density function (pdf) of the current magnitude and characterize the overall statistical profile of a trapped molecular state.

Specifically, the extracted features include the center of mass, standard deviation, peak height, coefficient of determination from Gaussian fitting, skewness, and kurtosis. Together, these six descriptors summarize the statistical properties of inhibitor-dependent current signatures while intentionally discarding temporal ordering and frequency information.

Wavelet features were extracted using the discrete wavelet transform (DWT) to jointly incorporate temporal and frequency-domain information. Unlike Fourier-based methods, which represent signals as sums of global sinusoidal components of varying frequency and implicitly assume stationarity, the DWT captures localized time–frequency representations that are well-suited for analyzing nonstationary signals containing transient fluctuations and within-event state transitions. The DWT is implemented through a cascade of high-pass and low-pass filtering operations followed by down sampling. The decomposition is defined by a scaling function *ø* (*t*) and its associated mother wavelet *ψ* (*t*) The mother wavelet is characterized by its order *p*, which determines the number of vanishing moments and ensures orthogonality to all polynomials of degree less than *p*. The filter length *F*, defined as the number of nonzero coefficients in the discrete filters described below, depends on the wavelet family and order p. The scaling function and associated mother wavelet satisfy the two-scale relations:

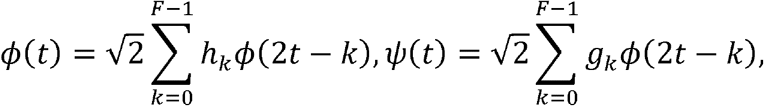

Where 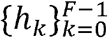 and 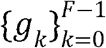 are the low-pass and high-pass filter coefficients, respectively, with the relation 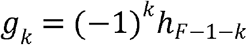

In the discrete setting, the sampled signal *x*[*n*] is interpreted via the scaling coefficients at the finest resolution level, i.e.a_0_[*n*] =*x*[*n*], and subsequent approximation and detail coefficients were obtained via a multiresolution filter-bank recursion over *L* levels:

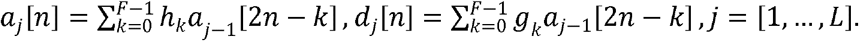

Here,*a*_*j*_[*n*] and d_*j*_[*n*] denote the approximation and detail coefficients at decomposition level _*j*_, respectively. The final decomposed coefficients contain *L* levels of detailed coefficients and one level of approximation coefficient:

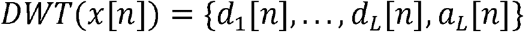

We used the PyWavelets framework and selected the Daubechies wavelet of order p = 4 (db-4), corresponding to a filter length *F* = 2*p*= 8 as the mother wavelet over *L* = 4 evels. This setting provides compact support and four vanishing moments, enabling efficient capture of abrupt current transitions while preserving low-frequency trends.

The resulting wavelet coefficients were subsequently transformed into fixed-length feature representations suitable for statistical learning. Specifically, four levels of wavelet decomposition produced one set of approximation coefficients a_4_[*n*] and four sets of detail coefficients {*d*_4_[*n*], *d*_3_[*n*], *d*_2_[*n*], *d*_1_[*n*]}, corresponding to progressively finer temporal scales. It is known that the approximation coefficients encode slowly varying, long-timescale current behavior, while the detail coefficients isolate higher-frequency fluctuations associated with short-lived transitions and transient dynamics within an event. To summarize the information contained in each coefficient set, ten statistical descriptors were computed from each level: mean, standard deviation, median, minimum, maximum, first quartile, third quartile, skewness, kurtosis, and mean absolute value. Concatenating these descriptors across all five coefficient sets results in a total of 50 wavelet-derived features per event.

To mitigate boundary artifacts arising from finite-length current traces, a symmetric signal extension was performed prior to the wavelet decomposition. For a discrete signal *x*[*n*] defined on *n*= 0,…, *N* − 1, samples outside the signal boundaries were generated as

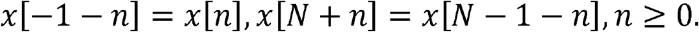

This enforced continuity of the signal and its local trends at the boundaries. It also prevented artificial discontinuities from introducing spurious high-frequency components into the wavelet coefficients, thereby ensuring that extracted features reflect intrinsic current fluctuations rather than edge effects.

Finally, baseline and wavelet features were concatenated to form a 56-dimensional feature vector for each event. To ensure comparable scaling across features and prevent dominance by variables with larger numerical ranges, all features were standardized using z-score normalization. Specifically, each feature *x* was transformed according to 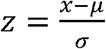,where *μ*and *σ* denote the mean and standard deviation of the feature computed over the training dataset, respectively. This normalization step ensures fair contributions of all features to downstream classification models and improves numerical stability during model training.

### Ensemble Classification

Supervised classification was performed using nine different models, including logistic regression, support vector machines (SVMs), random forests, XGBoost, LightGBM, k-nearest neighbors (k-NN), Gaussian naïve Bayes, multilayer perceptron (MLP), and quadratic discriminant analysis (QDA).

Let the full labeled dataset be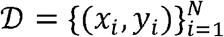, where *x*_*i*_ ∈ ℝ ^56^ denotes the feature vector of the *i*-th event and *y*_*i*_ ∈ {1,…,*C*} represents its class label. We partition ‘D into a training set *D*_*train*_ and a testing set *D*_*test*_ with an 80/20 split using stratified sampling, such that the class proportions are preserved in both splits, i.e.,

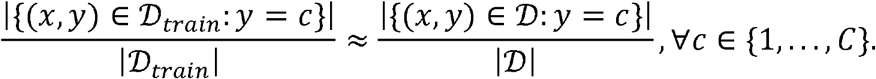

An analogously approach is used for *D*_*test*_.

Logistic regression learns a linear decision boundary in feature space and produces probabilistic class predictions via a softmax transformation, while SVMs construct a maximum-margin decision boundary, with nonlinear decision functions enabled through kernel mappings. Tree-based models (random forest, XGBoost, and LightGBM) learn ensembles of decision trees to capture nonlinear feature interactions, while gradient-boosted methods iteratively minimize a differentiable loss function via additive tree updates. The k-NN classifier assigns class labels based on the majority vote among the *k* nearest neighbors under a chosen distance metric. Gaussian naïve Bayes assumes conditional independence of features and class-conditional normal distributions. The MLP classifier learns nonlinear decision functions through stacked fully connected layers with nonlinear activations, while QDA models each class as a multivariate Gaussian distribution with class-specific covariance matrices.

To improve robustness and reduce model-specific bias, ensemble classifiers were constructed using hard voting, soft voting, and stacking. In hard voting, the final predicted label 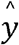 is determined by a majority vote:

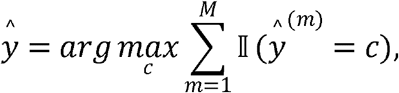

Where 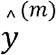 is the prediction of the *m*-th base classifier and *M*= 9 is the number of classifiers used. In soft voting, class probabilities are averaged across classifiers:

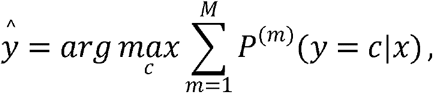

where *p*(^m)^ (y = c|x) denotes the predicted probability from the *m*-th model. In stacking, the outputs of base classifiers are concatenated and used as inputs to a meta-classifier *f*_*meta*_, resulting in the final prediction:

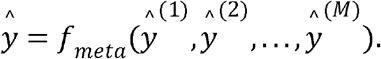

Ensemble strategies leverage complementary strengths of individual classifiers to achieve more stable and accurate predictions.

### Mixture data prediction

For mixture-data experiments including multiple inhibitors, individual trapping events frequently consisted of both apo-like and inhibitor-bound segments without clear escape peaks separating molecular states. To identify distinct segments within a single event, we applied a rolling-average filter with a window length of 0.1s to the residual current trace. For a discrete signal *x*[*n*], the rolling average 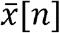 is computed as

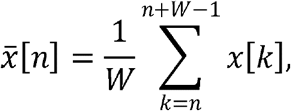

where *W* corresponds to 0.1s of data (1,000 samples at a 10 kHz sampling rate). An amplitude threshold of 40% residual current *I*_*res*_ is applied to the rolling-average signal 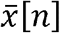. Crossings of this threshold were used to partition a single event into multiple contiguous segments. Each resulting segment was subsequently treated as a sample for subsequent analysis. Baseline and wavelet features are extracted separately from each segment, and the trained classifiers and ensemble models were used to predict the inhibitor identity for each segment.

## Author contributions

Ly N. designed and performed experiments, analyzed data, and wrote the manuscript, Y.W. developed computational algorithms, analyzed data, and wrote the manuscript. J.C.F. performed experiments and data analysis, D.D. contributed to data processing, Lan N. and B.W. assisted in current recording experiments. O.M. and M.C. supervised the project and edited the manuscript.

## Acknowledgments

The research in the Chen lab was supported by the US National Institutes of Health grant R01AI156187 and R01GM159415.

## Notes

### Competing Interest Statement

The authors have declared no competing interest.

## References

1. Bhullar, K. S. et al. Kinase-targeted cancer therapies: progress, challenges and future directions. Mol. Cancer 17, 48 (2018).

2. Druker, B. J. et al. Activity of a Specific Inhibitor of the BCR-ABL Tyrosine Kinase in the Blast Crisis of Chronic Myeloid Leukemia and Acute Lymphoblastic Leukemia with the Philadelphia Chromosome. N. Engl. J. Med. 344, 1038–1042 (2001).

3. Knapp, S. New opportunities for kinase drug repurposing and target discovery. Br. J. Cancer 118, 936–937 (2018).

4. Liebl, E. C. et al. Dosage-Sensitive, Reciprocal Genetic Interactions between the Abl Tyrosine Kinase and the Putative GEF trio Reveal trio’s Role in Axon Pathfinding. Neuron 26, 107–118 (2000).

5. Lebouvier, T. et al. The Microtubule-Associated Protein Tau is Also Phosphorylated on Tyrosine. J. Alzheimer’s Dis. 18, 1–9 (2009).

6. Arter, C., Trask, L., Ward, S., Yeoh, S. & Bayliss, R. Structural features of the protein kinase domain and targeted binding by small-molecule inhibitors. J. Biol. Chem. 298, (2022).

7. Dar, A. C. & Shokat, K. M. The Evolution of Protein Kinase Inhibitors from Antagonists to Agonists of Cellular Signaling. Annu. Rev. Biochem. 80, 769–795 (2011).

8. Dang, X.-W. et al. Recent advances of small-molecule c-Src inhibitors for potential therapeutic utilities. Bioorganic Chem. 142, 106934 (2024).

9. Capdeville, R., Buchdunger, E., Zimmermann, J. & Matter, A. Glivec (STI571, imatinib), a rationally developed, targeted anticancer drug. Nat. Rev. Drug Discov. 1, 493–502 (2002).

10. Roskoski, R. Properties of FDA-approved small molecule protein kinase inhibitors: A 2026 update. Pharmacol. Res. 224, 108107 (2026).

11. Elgawish, M. S., Almatary, A. M., Zaitone, S. A. & Salem, M. S. H. Leveraging artificial intelligence and machine learning in kinase inhibitor development: advances, challenges, and future prospects. RSC Med. Chem. 16, 4698–4720 (2025).

12. Roskoski, R. Classification of small molecule protein kinase inhibitors based upon the structures of their drug-enzyme complexes. Pharmacol. Res. 103, 26–48 (2016).

13. Meharena, H. S. et al. Deciphering the Structural Basis of Eukaryotic Protein Kinase Regulation. PLOS Biol. 11, e1001680 (2013).

14. Mian, A. A. et al. The gatekeeper mutation T315I confers resistance against small molecules by increasing or restoring the ABL-kinase activity accompanied by aberrant transphosphorylation of endogenous BCR, even in loss-of-function mutants of BCR/ABL. Leukemia 23, 1614–1621 (2009).

15. O’Hare, T. et al. AP24534, a Pan-BCR-ABL Inhibitor for Chronic Myeloid Leukemia, Potently Inhibits the T315I Mutant and Overcomes Mutation-Based Resistance. Cancer Cell 16, 401–412 (2009).

16. Modi, V. & Dunbrack, R. L., Jr. Kincore: a web resource for structural classification of protein kinases and their inhibitors. Nucleic Acids Res. 50, D654–D664 (2022).

17. Joseph, R. E. et al. Differential impact of BTK active site inhibitors on the conformational state of full-length BTK. eLife 9, e60470 (2020).

18. Constantine, K. L. et al. Multiple and Single Binding Modes of Fragment-Like Kinase Inhibitors Revealed by Molecular Modeling, Residue Type-Selective Protonation, and Nuclear Overhauser Effects. J. Med. Chem. 51, 6225–6229 (2008).

19. Anderson, J. W. et al. Conformation selection by ATP-competitive inhibitors and allosteric communication in ERK2. eLife 12, RP91507 (2024).

20. Simard, J. R. et al. Fluorophore Labeling of the Glycine-Rich Loop as a Method of Identifying Inhibitors That Bind to Active and Inactive Kinase Conformations. J. Am. Chem. Soc. 132, 4152–4160 (2010).

21. Straathof, S., Di Muccio, G. & Maglia, G. Nanopores with an Engineered Selective Entropic Gate Detect Proteins at Nanomolar Concentration in Complex Biological Sample. J. Am. Chem. Soc. 147, 15050–15065 (2025).

22. Zhang, X. et al. Specific Detection of Proteins by a Nanobody-Functionalized Nanopore Sensor. ACS Nano 17, 9167–9177 (2023).

23. Motone, K. et al. Multi-pass, single-molecule nanopore reading of long protein strands. Nature 633, 662–669 (2024).

24. Bonini, A. et al. Single-molecule identification of full-length proteins with single-amino-acid resolution using nanopores. 2026.01.08.698148 Preprint at 10.64898/2026.01.08.698148 (2026).

25. Fahie, M. A. & Chen, M. Electrostatic Interactions between OmpG Nanopore and Analyte Protein Surface Can Distinguish between Glycosylated Isoforms. J. Phys. Chem. B 119, 10198–10206 (2015).

26. Soni, N. et al. Full-length protein classification via cysteine fingerprinting in solid-state nanopores. Nat. Nanotechnol. 20, 1482–1490 (2025).

27. Jo, J., Kim, J.-S., Hwang, S.Jeong, K.-B. & Chi, S.-W. Nanopore Discrimination of Protein–Small-Molecule Drug Complexes at Near-Atomic Resolution. ACS Nano 20, 13624–13635 (2026).

28. Li, X., Lee, K. H., Shorkey, S., Chen, J. & Chen, M. Different Anomeric Sugar Bound States of Maltose Binding Protein Resolved by a Cytolysin A Nanopore Tweezer. ACS Nano 14, 1727–1737 (2020).

29. Soskine, M., Biesemans, A. & Maglia, G. Single-Molecule Analyte Recognition with ClyA Nanopores Equipped with Internal Protein Adaptors. J. Am. Chem. Soc. 137, 5793–5797 (2015).

30. Nova, I. C. et al. Detection of phosphorylation post-translational modifications along single peptides with nanopores. Nat. Biotechnol. 42, 710–714 (2024).

31. Li, S. et al. T232K/K238Q Aerolysin Nanopore for Mapping Adjacent Phosphorylation Sites of a Single Tau Peptide. Small Methods 4, 2000014 (2020).

32. Cao, C. et al. Deep Learning-Assisted Single-Molecule Detection of Protein Post-translational Modifications with a Biological Nanopore. ACS Nano 18, 1504–1515 (2024).

33. Lan, W.-H., He, H., Bayley, H. & Qing, Y. Location of Phosphorylation Sites within Long Polypeptide Chains by Binder-Assisted Nanopore Detection. J. Am. Chem. Soc. 146, 24265–24270 (2024).

34. Sexton, L. T. et al. Resistive-Pulse Studies of Proteins and Protein/Antibody Complexes Using a Conical Nanotube Sensor. J. Am. Chem. Soc. 129, 13144–13152 (2007).

35. Thakur, A. K. & Movileanu, L. Real-time measurement of protein–protein interactions at single-molecule resolution using a biological nanopore. Nat. Biotechnol. 37, 96–101 (2019).

36. Sun, J., Skanata, A. & Movileanu, L. Single-Molecule Observation of Competitive Protein– Protein Interactions Utilizing a Nanopore. ACS Nano 19, 1103–1115 (2024).

37. Sarthak, K., Van Meervelt, V., Soskine, M., Maglia, G. & Aksimentiev, A. Engineering a Biological Nanopore for Monitoring Protein Dynamics and Conformational Changes at the Single-Molecule Level. ACS Nano 20, 18848–18862 (2026).

38. Kavetsky, K., Hong, S., Lin, C.-Y., Yang, R. & Drndić, M. Uncovering Hidden Protein Conformations with High Bandwidth Nanopore Measurements. Nano Lett. 26, 2709–2716 (2026).

39. Li, F., Fahie, M. A., Gilliam, K. M., Pham, R. & Chen, M. Mapping the conformational energy landscape of Abl kinase using ClyA nanopore tweezers. Nat. Commun. 13, 3541 (2022).

40. Shorkey, S. A. et al. Tracking flaviviral protease conformational dynamics by tuning single-molecule nanopore tweezers. Biophys. J. 124, 145–157 (2025).

41. Soskine, M., Biesemans, A., De Maeyer, M. & Maglia, G. Tuning the Size and Properties of ClyA Nanopores Assisted by Directed Evolution. J. Am. Chem. Soc. 135, 13456–13463 (2013).

42. Jeong, K.-B. et al. Single-molecule fingerprinting of protein-drug interaction using a funneled biological nanopore. Nat. Commun. 14, 1461 (2023).

43. Straathof, S. et al. Protein Sizing with 15 nm Conical Biological Nanopore YaxAB. ACS Nano 17, 13685–13699 (2023).

44. Galenkamp, N. S. & Maglia, G. Single-Molecule Sampling of Dihydrofolate Reductase Shows Kinetic Pauses and an Endosteric Effect Linked to Catalysis. ACS Catal. 12, 1228–1236 (2022).

45. Li, F., Foster, J. C., Nguyen, L. & Chen, M. Selective Identification of Allosteric Inhibitors and Co-Drug Combinations Targeting Kinases by Using a Nanopore Tweezer Approach. ACS Nano 19, 34617–34627 (2025).

46. Tokarski, J. S. et al. The Structure of Dasatinib (BMS-354825) Bound to Activated ABL Kinase Domain Elucidates Its Inhibitory Activity against Imatinib-Resistant ABL Mutants. Cancer Res. 66, 5790–5797 (2006).

47. Nagar, B. et al. Structural Basis for the Autoinhibition of c-Abl Tyrosine Kinase. Cell 112, 859–871 (2003).

48. Cowan-Jacob, S. W. et al. Structural biology contributions to the discovery of drugs to treat chronic myelogenous leukaemia. Acta Crystallogr. D Biol. Crystallogr. 63, 80–93 (2007).

49. Zhou, T. et al. Structural Mechanism of the Pan-BCR-ABL Inhibitor Ponatinib (AP24534): Lessons for Overcoming Kinase Inhibitor Resistance. Chem. Biol. Drug Des. 77, 1–11 (2011).

50. Weisberg, E. et al. Characterization of AMN107, a selective inhibitor of native and mutant Bcr-Abl. Cancer Cell 7, 129–141 (2005).

51. Hochhaus, A. et al. Nilotinib is associated with a reduced incidence of BCR-ABL mutations vs imatinib in patients with newly diagnosed chronic myeloid leukemia in chronic phase. Blood 121, 3703–3708 (2013).

52. Levinson, N. M. & Boxer, S. G. Structural and Spectroscopic Analysis of the Kinase Inhibitor Bosutinib and an Isomer of Bosutinib Binding to the Abl Tyrosine Kinase Domain. PLOS ONE 7, e29828 (2012).

53. Knowles, P. P. et al. Structure and Chemical Inhibition of the RET Tyrosine Kinase Domain*. J. Biol. Chem. 281, 33577–33587 (2006).

54. Musumeci, F., Schenone, S., Brullo, C. & Botta, M. An Update On Dual Src/Abl Inhibitors. Future Med. Chem. 4, 799–822 (2012).

55. Hennequin, L. F. et al. N-(5-Chloro-1,3-benzodioxol-4-yl)-7-[2-(4-methylpiperazin-1-yl)ethoxy]-5-(tetrahydro-2H-pyran-4-yloxy)quinazolin-4-amine, a Novel, Highly Selective, Orally Available, Dual-Specific c-Src/Abl Kinase Inhibitor. J. Med. Chem. 49, 6465–6488 (2006).

56. Gucalp, A. et al. Phase II trial of saracatinib (AZD0530), an oral src-inhibitor for the treatment of patients with hormone receptor negative metastatic breast cancer. Clin. Breast Cancer 11, 306–311 (2011).

57. Xie, T., Saleh, T., Rossi, P. & Kalodimos, C. G. Conformational states dynamically populated by a kinase determine its function. Science 370, eabc2754 (2020).

58. Gibbons, D. L., Pricl, S., Kantarjian, H., Cortes, J. & Quintás-Cardama, A. The rise and fall of gatekeeper mutations? The BCR-ABL1 T315I paradigm. Cancer 118, 293–299 (2012).

59. Branford, S. et al. Detection of BCR-ABL mutations in patients with CML treated with imatinib is virtually always accompanied by clinical resistance, and mutations in the ATP phosphate-binding loop (P-loop) are associated with a poor prognosis. Blood 102, 276–283 (2003).

60. von Bubnoff, N. et al. Bcr-Abl resistance screening predicts a limited spectrum of point mutations to be associated with clinical resistance to the Abl kinase inhibitor nilotinib (AMN107). Blood 108, 1328–1333 (2006).

61. O’Hare, T., Eide, C. A. & Deininger, M. W. N. Bcr-Abl kinase domain mutations, drug resistance, and the road to a cure for chronic myeloid leukemia. Blood 110, 2242–2249 (2007).

62. Lyczek, A. et al. Mutation in Abl kinase with altered drug-binding kinetics indicates a novel mechanism of imatinib resistance. Proc. Natl. Acad. Sci. 118, e2111451118 (2021).

