## Supplementary Information for "Subtyping Abl Kinase Inhibitor Binding Modes with Machine-Learning-Enabled Super Resolution Single-Molecule Nanopore Tweezers"

**Table of Contents**

List of Abbreviations

Material

Figure S1. Conformational states of the Abl kinase domain

Figure S2. Representative current traces Apo Abl-WT

Figure S3. Representative current traces of dasatinib-dosed Abl WT

Figure S4. Representative current traces of vandatenib-dosed Abl WT

Figure S5. Representative current traces of imatinib-dosed Abl WT

Figure S6. Representative current traces of nilotinib-dosed Abl WT

Figure S7. Representative current traces of ponatinib-dosed Abl WT

Figure S8. Representative current traces of bosutinib-dosed Abl WT

Figure S9. Representative current traces of saracatinib-dosed Abl WT

Figure S10. Representative current traces Apo Abl T315I

Figure S11. Representative current traces Apo Abl E255V

Figure S12. Representative current traces of dasatinib-dosed Abl T315I

Figure S13. Representative current traces of vandatenib-dosed Abl T315I

Figure S14. Representative current traces of imatinib-dosed Abl T315I

Figure S15. Representative current traces of nilotinib-dosed Abl T315I

Figure S16. Representative current traces of ponatinib-dosed Abl T315I

Figure S17. Representative current traces of bosutinib-dosed Abl T315I

Figure S18. Representative current traces of saracatinib-dosed Abl T315I

Figure S19. Representative current traces of dasatinib-dosed Abl E255V

Figure S20. Representative current traces of vandatenib-dosed Abl E255V

Figure S21. Representative current traces of imatinib-dosed Abl E255V

Figure S22. Representative current traces of nilotinib-dosed Abl E255V

Figure S23. Representative current traces of ponatinib-dosed Abl E255V

Figure S24. Representative current traces of bosutinib-dosed Abl E255V

Figure S25. Representative current traces of saracatinib-dosed Abl E255V

Figure S26 Example inhibitor bound/unbound analysis traces.

Figure S27 Representative predicted results from inhibitor mixture A dosed to Abl WT
Figure S28 Representative predicted results from inhibitor mixture B1 dosed to Abl WT
Figure S29 Representative predicted results from inhibitor mixture B2 dosed to Abl WT
Table S1 Quantitative analysis results for inhibitor mixtures dosed to Abl WT
Table S2 Primers used for generating Abl variants by site-directed mutagenesis
References

**List of Abbreviations**

| Abbreviation | Definition |
|---|---|
| Bos | bosutinib |
| Das | dasatinib |
| Ima | imatinib |
| Nil | nilotinib |
| Pon | ponatinib |
| Sar | saracatinib |
| Van | vandetanib |

### Materials

Bosutinib (4-[(2,4-dichloro-5-methoxyphenyl)amino]-6-methoxy-7-[3-(4-methylpiperazin-1-yl)propoxy]quinoline-3-carbonitrile), dasatinib (N-(2-chloro-6-methylphenyl)-2-[[6-[4-(2-hydroxyethyl)-1-piperazinyl]-2-methyl-4-pyrimidinyl]amino]-5-thiazole carboxamide monohydrate), imatinib (4-[(4-methylpiperazin-1-yl)methyl]-N-(4-methyl-3-{[4-(pyridin-3-yl)pyrimidin-2-yl]amino}phenyl)), nilotinib (4-methyl-N-[3-(4-methyl-1H-imidazol-1-yl)- 5-(trifluoromethyl)phenyl]-3- [(4-pyridin-3-ylpyrimidin-2-yl) amino]benzamide), ponatinib (3-(2-Imidazo[1,2-b]pyridazin-3-ylethynyl)-4-methyl-N-[4-[(4-methylpiperazin-1-yl)methyl]-3-(trifluoromethyl)phenyl]benzamide), saracatinib (N-(5-chloro-1,3-benzodioxol-4-yl)-7-[2-(4-methyl-1-piperazinyl)ethoxy]-5-[(tetrahydro-2H-pyran-4-yl)oxy]-4-quinazolinamine), vandetanib (N-(4-bromo-2-fluorophenyl)-6-methoxy-7-[(1-methylpiperidin-4-yl)methoxy]quinazolin-4-amine) were purchased from MedChem Express.

Disodium phosphate ($Na_2HPO_4$), dimethylsulfoxide (DMSO), glycerol, monopotassium phosphate ($KH_2PO_4$), potassium chloride (KCl), sodium chloride (NaCl), and labware were purchased from Fisher Scientific. Isopropyl-β-D-thiogalactopyranoside (IPTG) and n-Dodecyl-B-D-maltoside (DDM) were purchased from GoldBio. HisPur™ Ni-NTA resin was purchased from Thermo Fisher Scientific. HW55s resin was purchased from Tosoh Bioscience. 1,2-Diphytanoyl-sn-glycerol-3-phosphocholine (DPhPC) lipids were purchased from Avanti Polar Lipids. Silver for electrodes was purchased from Alfa Aesar. All other reagents were purchased from Research Products International unless otherwise stated.

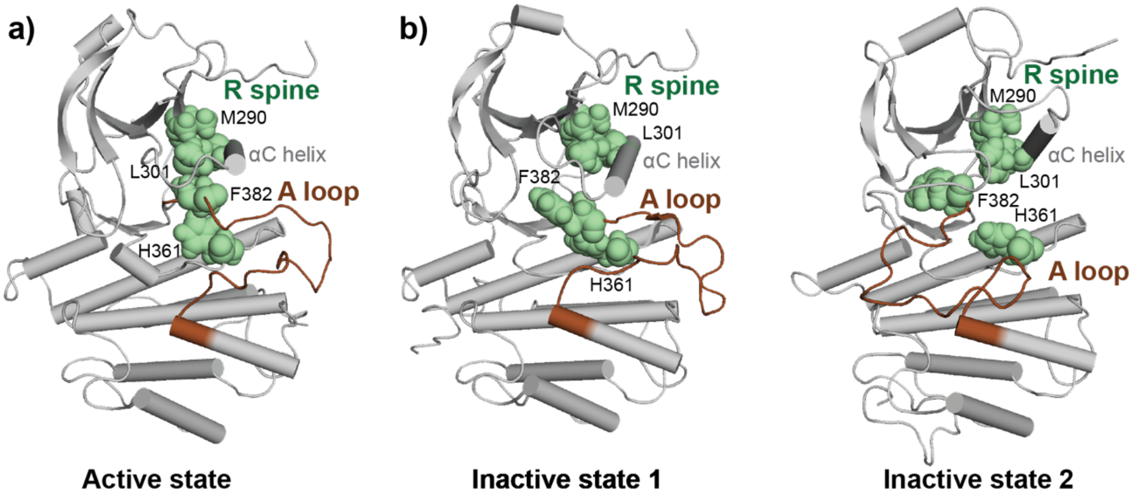


**Figure S1.** Conformational states of the Abl kinase domain with the R-spine, made up by M290, L301, H361 and F382 residues, highlighted in green and the A-loop highlighted in brown. **(a)** Assembled R-spine in the active state of Abl kinase (PDB: 6XR6). **(b)** Dismantled R-spine in the inactive states of Abl kinase (PDB: 6XR7 and 6XRG).[1]

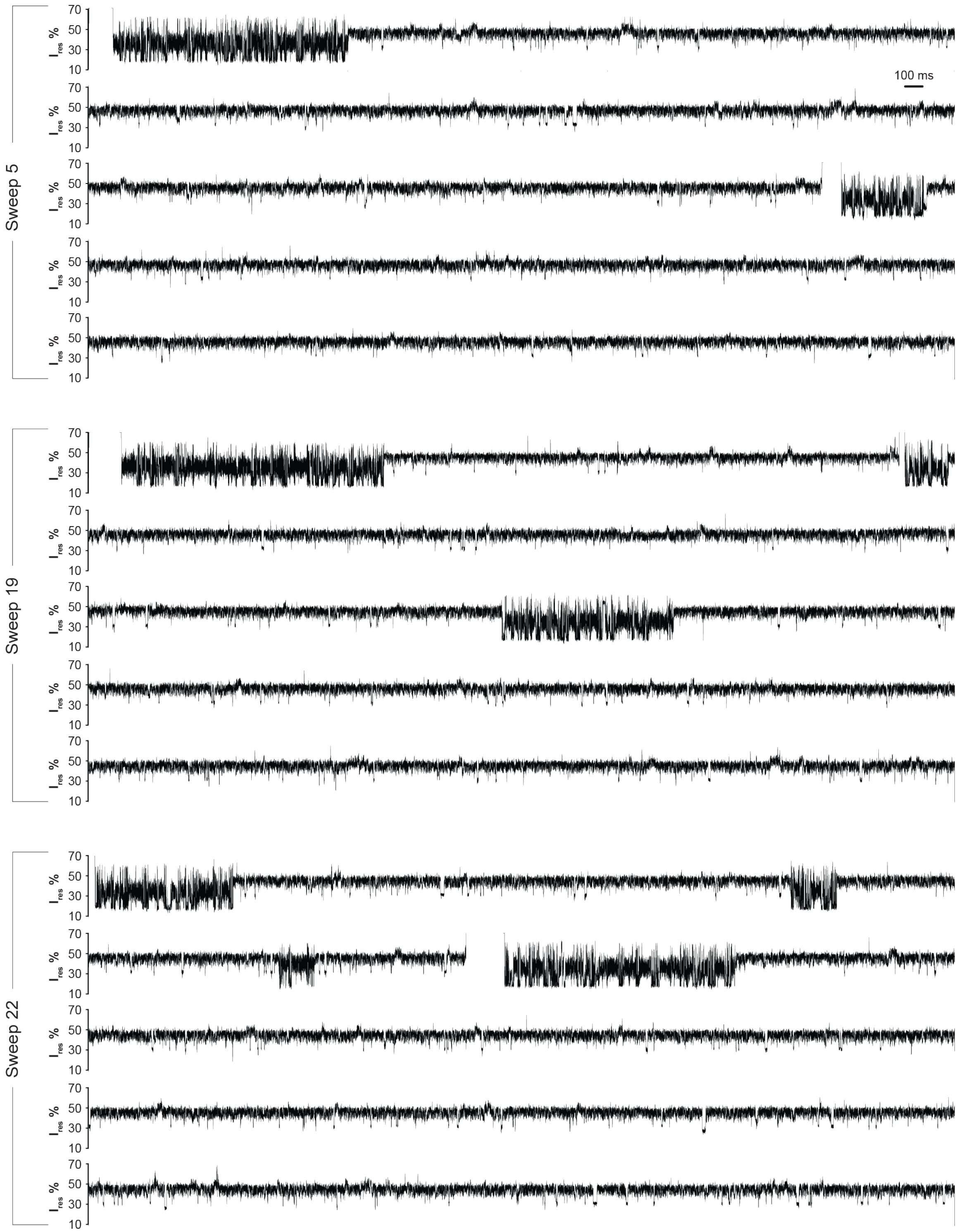


**Figure S2.** Three representative current traces of Apo Abl-WT recorded using the sweep protocol. Each trace spans 25s and was obtained at a trapping voltage of −80 mV. The $I_{res}$% range (10–70%) is shown to highlight S1/S2 signal patterns. Open-pore current and the zero baseline fall outside the displayed $I_{res}$ range and therefore appear as blank regions. Recordings were performed in 100 mM Tris-HCl (pH 7.5), 150 mM NaCl, 10 mM $MgCl_2$, and 1 mM DTT.

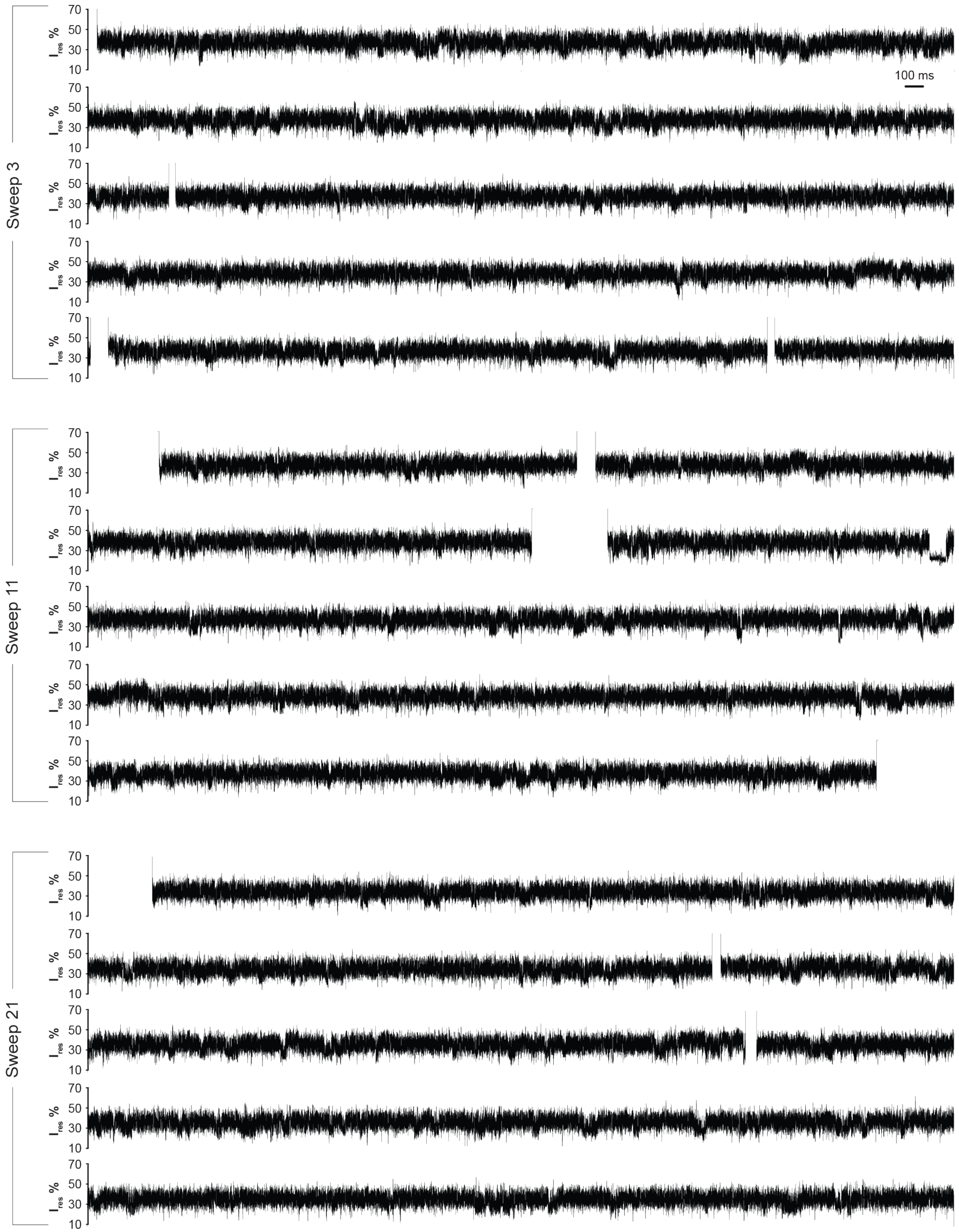


**Figure S3.** Three representative current traces of dasatinib-dosed Abl-WT recorded using the sweep protocol. Each trace spans 25s and was obtained at a trapping voltage of −80 mV. The $I_{res}$% range (10–70%) is shown to highlight S1/S2 signal patterns. Open-pore current and the zero baseline fall outside the displayed $I_{res}$ range and therefore appear as blank regions. Recordings were performed in 100 mM Tris-HCl (pH 7.5), 150 mM NaCl, 10 mM $MgCl_2$, and 1 mM DTT, supplemented with 150 nM dasatinib at 0.15% (v/v) DMSO (final concentrations).

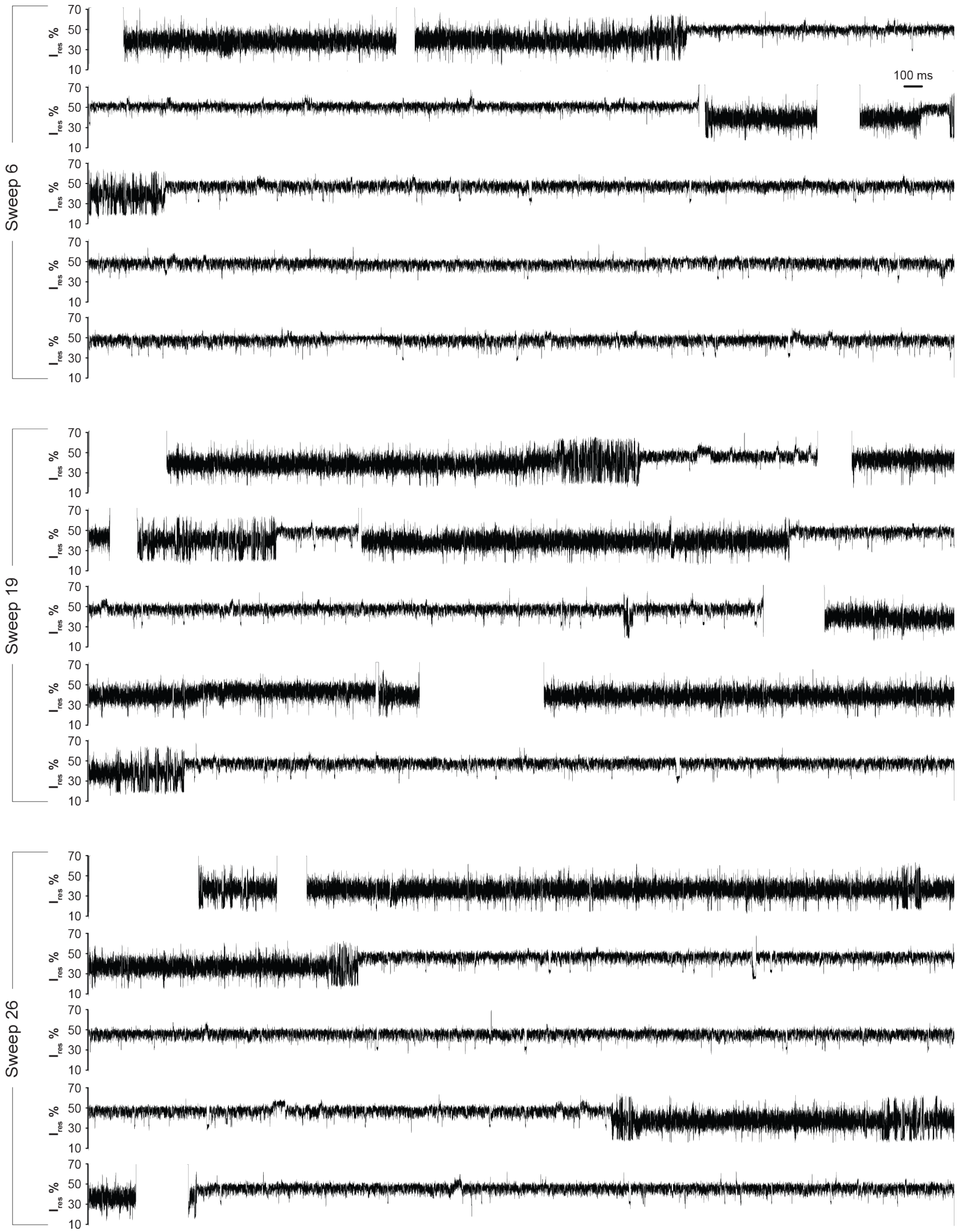


**Figure S4.** Three representative current traces of vandatenib-dosed Abl-WT recorded using the sweep protocol. Each trace spans 25 s and was obtained at a trapping voltage of −80 mV. The $I_{res}$% range (10–70%) is shown to highlight S1/S2 signal patterns. Open-pore current and the zero baseline fall outside the displayed $I_{res}$ range and therefore appear as blank regions. Recordings were performed in 100 mM Tris-HCl (pH 7.5), 150 mM NaCl, 10 mM $MgCl_2$, and 1 mM DTT, supplemented with 150 nM vandatenib at 0.15% (v/v) DMSO (final concentrations).

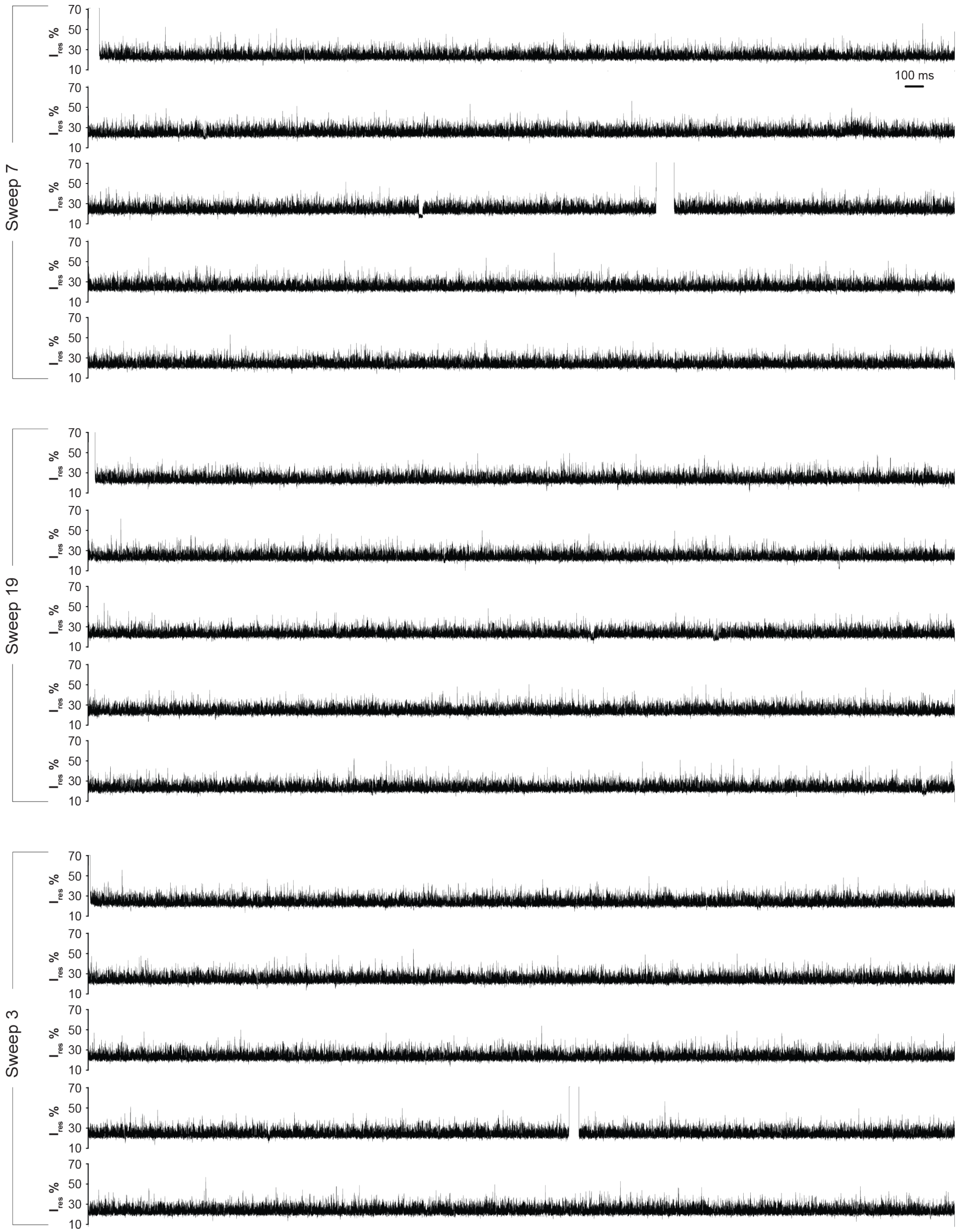


**Figure S5.** Three representative current traces of imatinib-dosed Abl-WT recorded using the sweep protocol. Each trace spans 25 s and was obtained at a trapping voltage of −80 mV. The $I_{res}$% range (10–70%) is shown to highlight S1/S2 signal patterns. Open-pore current and the zero baseline fall outside the displayed $I_{res}$ range and therefore appear as blank regions. Recordings were performed in 100 mM Tris-HCl (pH 7.5), 150 mM NaCl, 10 mM $MgCl_2$, and 1 mM DTT, supplemented with 150 nM imatinib at 0.15% (v/v) DMSO (final concentrations).

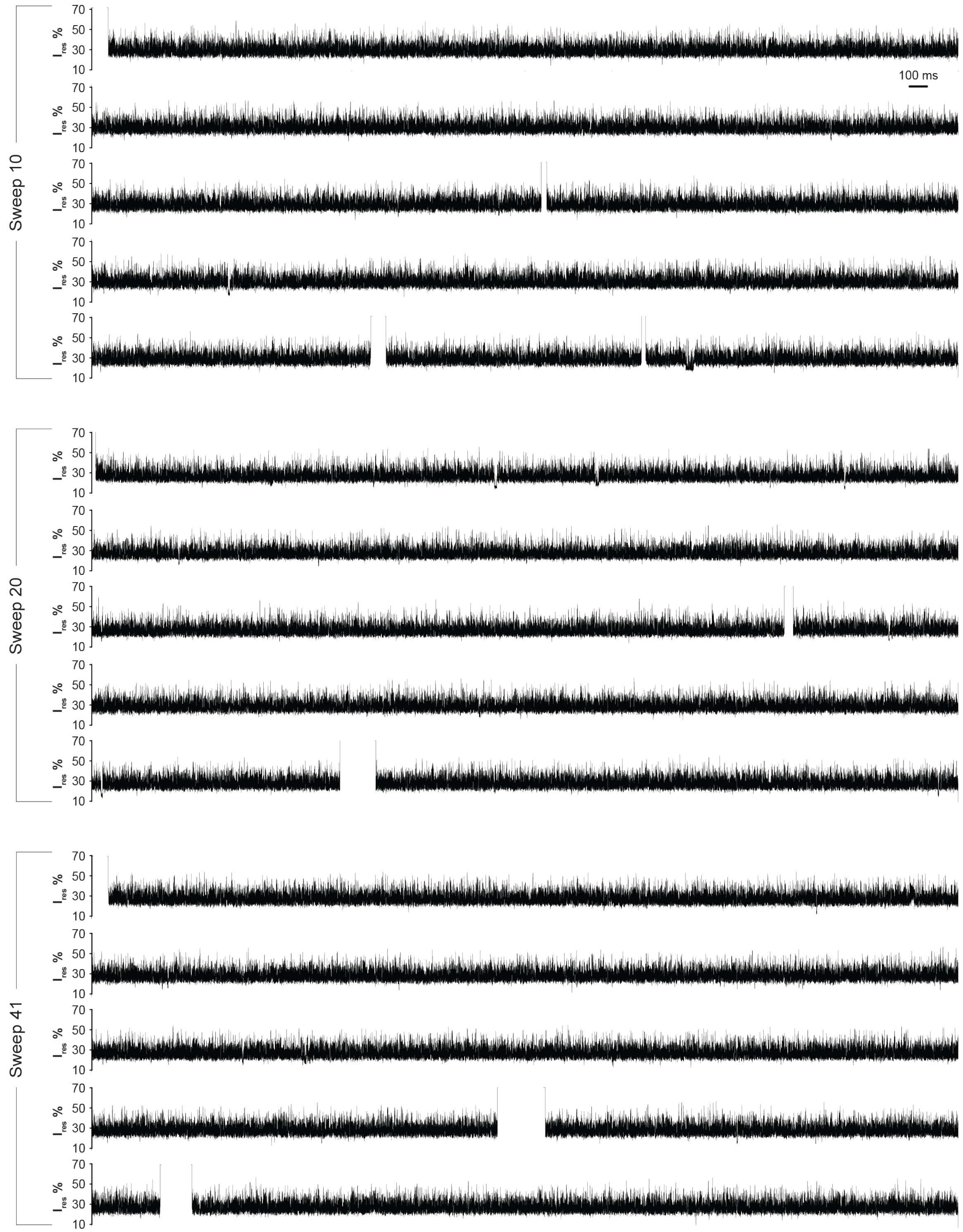


**Figure S6.** Three representative current traces of nilotinib-dosed Abl-WT recorded using the sweep protocol. Each trace spans 25 s and was obtained at a trapping voltage of −80 mV. The $I_{res}$% range (10–70%) is shown to highlight S1/S2 signal patterns. Open-pore current and the zero baseline fall outside the displayed $I_{res}$ range and therefore appear as blank regions. Recordings were performed in 100 mM Tris-HCl (pH 7.5), 150 mM NaCl, 10 mM $MgCl_2$, and 1 mM DTT, supplemented with 150 nM nilotinib at 0.15% (v/v) DMSO (final concentrations).

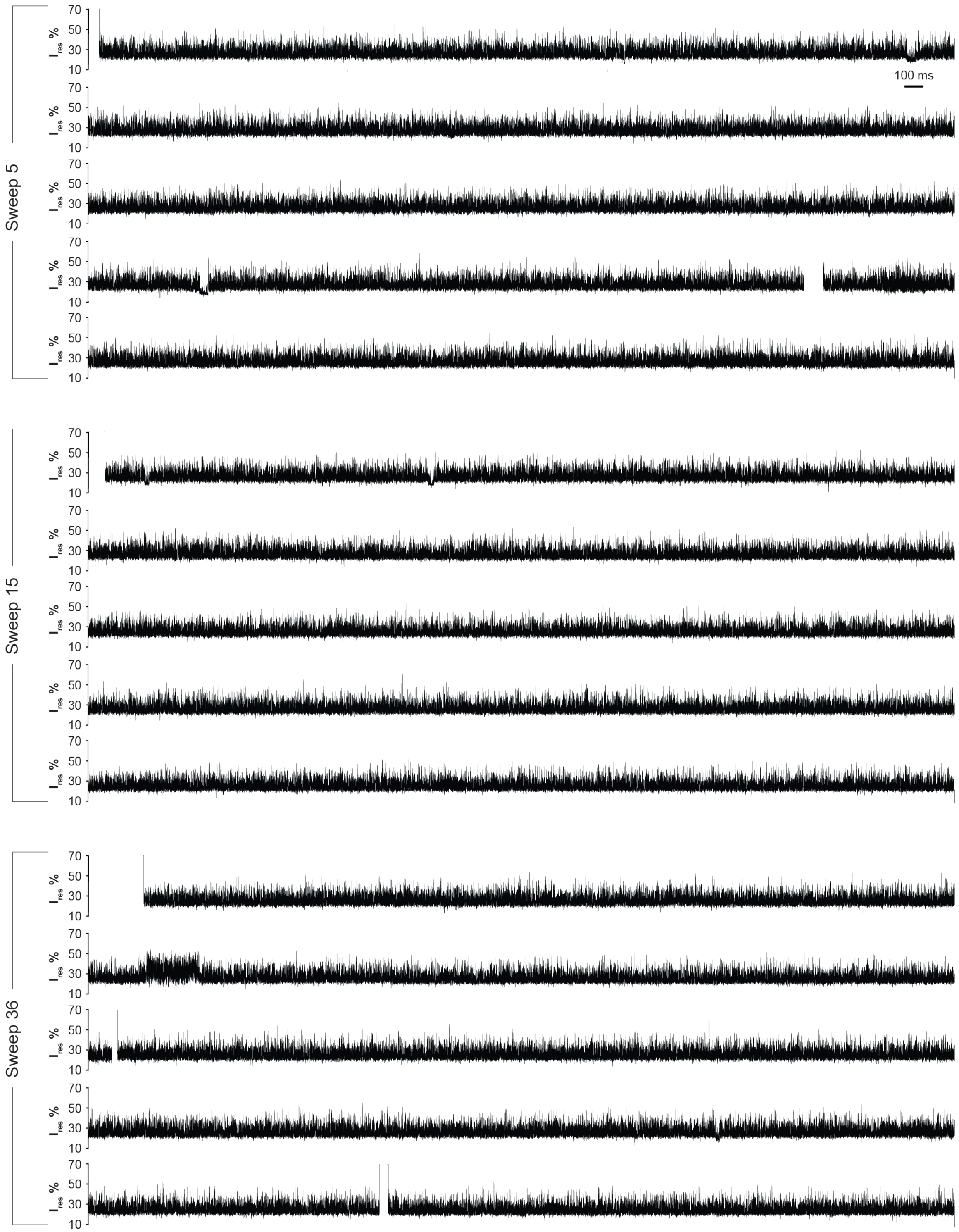


**Figure S7.** Three representative current traces of ponatinib-dosed Abl-WT recorded using the sweep protocol. Each trace spans 25 s and was obtained at a trapping voltage of −80 mV. The $I_{res}$% range (10–70%) is shown to highlight S1/S2 signal patterns. Open-pore current and the zero baseline fall outside the displayed $I_{res}$ range and therefore appear as blank regions. Recordings were performed in 100 mM Tris-HCl (pH 7.5), 150 mM NaCl, 10 mM $MgCl_2$, and 1 mM DTT, supplemented with 150 nM ponatinib at 0.15% (v/v) DMSO (final concentrations).

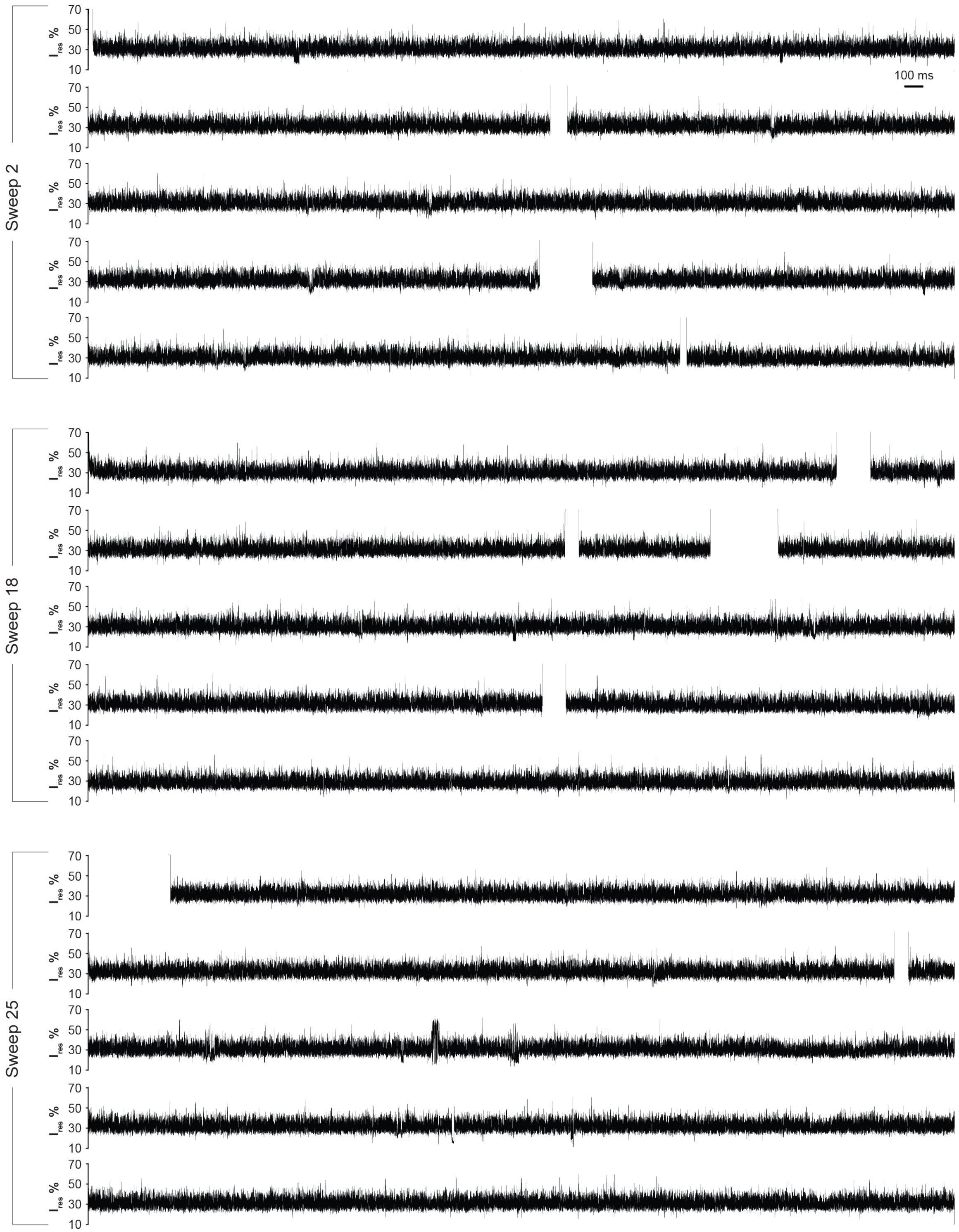


**Figure S8.** Three representative current traces of bosutinib-dosed Abl-WT recorded using the sweep protocol. Each trace spans 25 s and was obtained at a trapping voltage of −80 mV. The $I_{res}$% range (10–70%) is shown to highlight S1/S2 signal patterns. Open-pore current and the zero baseline fall outside the displayed $I_{res}$ range and therefore appear as blank regions. Recordings were performed in 100 mM Tris-HCl (pH 7.5), 150 mM NaCl, 10 mM $MgCl_2$, and 1 mM DTT, supplemented with 150 nM bosutinib at 0.15% (v/v) DMSO (final concentrations).

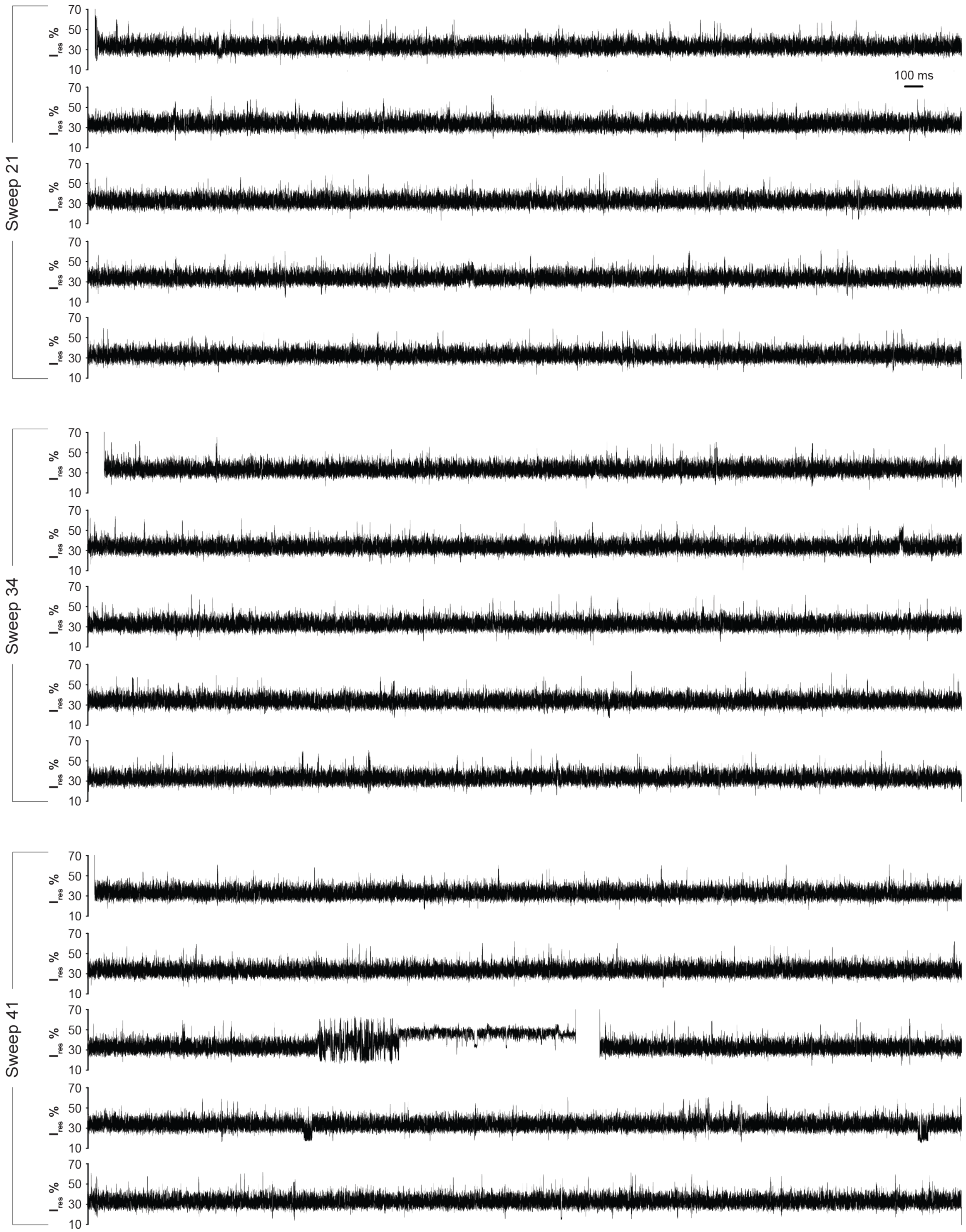


**Figure S9.** Three representative current traces of saracatinib-dosed Abl-WT recorded using the sweep protocol. Each trace spans 25 s and was obtained at a trapping voltage of −80 mV. The $I_{res}$% range (10–70%) is shown to highlight S1/S2 signal patterns. Open-pore current and the zero baseline fall outside the displayed $I_{res}$ range and therefore appear as blank regions. Recordings were performed in 100 mM Tris-HCl (pH 7.5), 150 mM NaCl, 10 mM $MgCl_2$, and 1 mM DTT, supplemented with 150 nM saracatinib at 0.15% (v/v) DMSO (final concentrations).

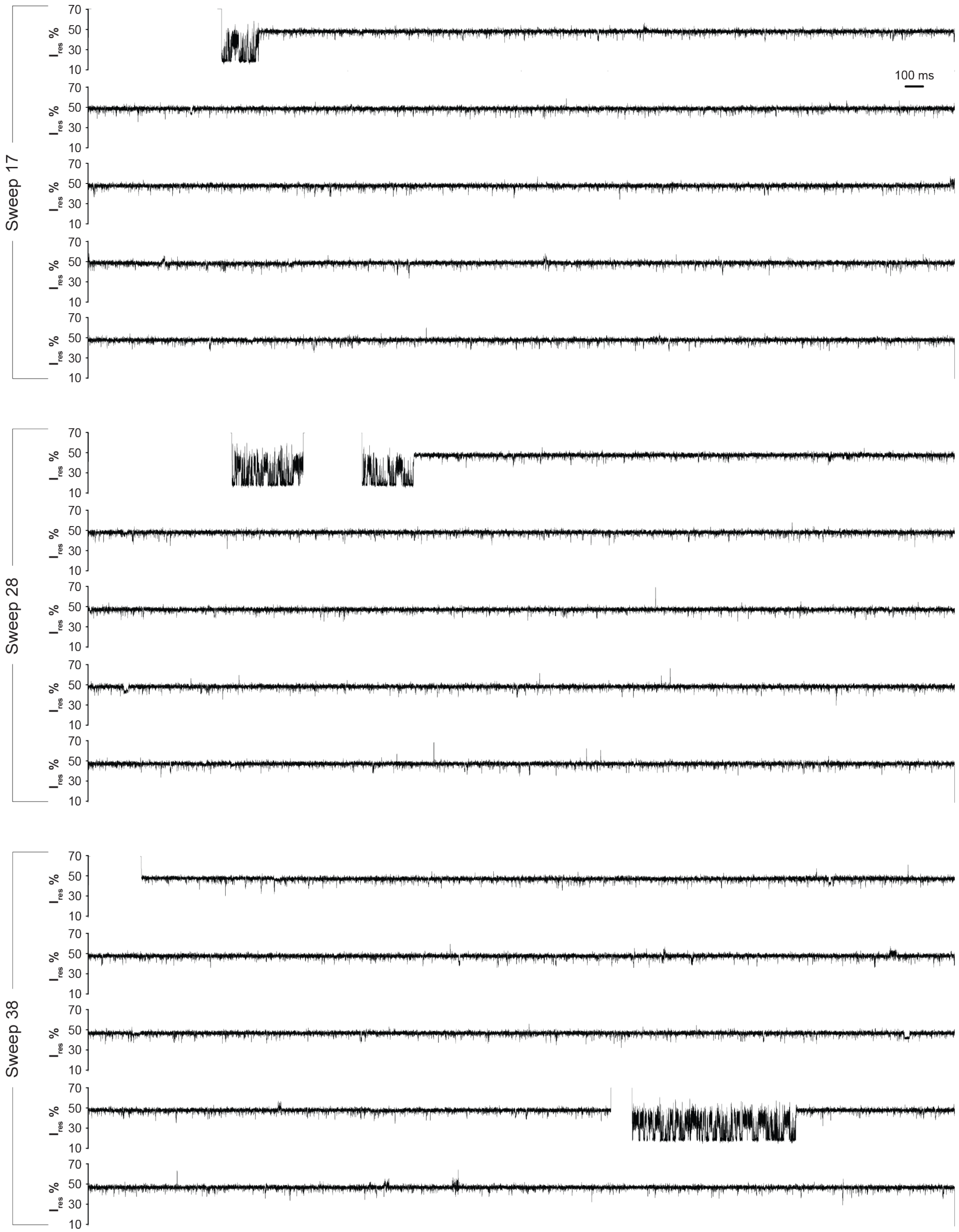


**Figure S10.** Three representative current traces Apo Abl T315I recorded using the sweep protocol. Each trace spans 25 s and was obtained at a trapping voltage of −80 mV. The $I_{res}$% range (10–70%) is shown to highlight S1/S2 signal patterns. Open-pore current and the zero baseline fall outside the displayed $I_{res}$ range and therefore appear as blank regions. Recordings were performed in 100 mM Tris-HCl (pH 7.5), 150 mM NaCl, 10 mM $MgCl_2$, and 1 mM DTT.

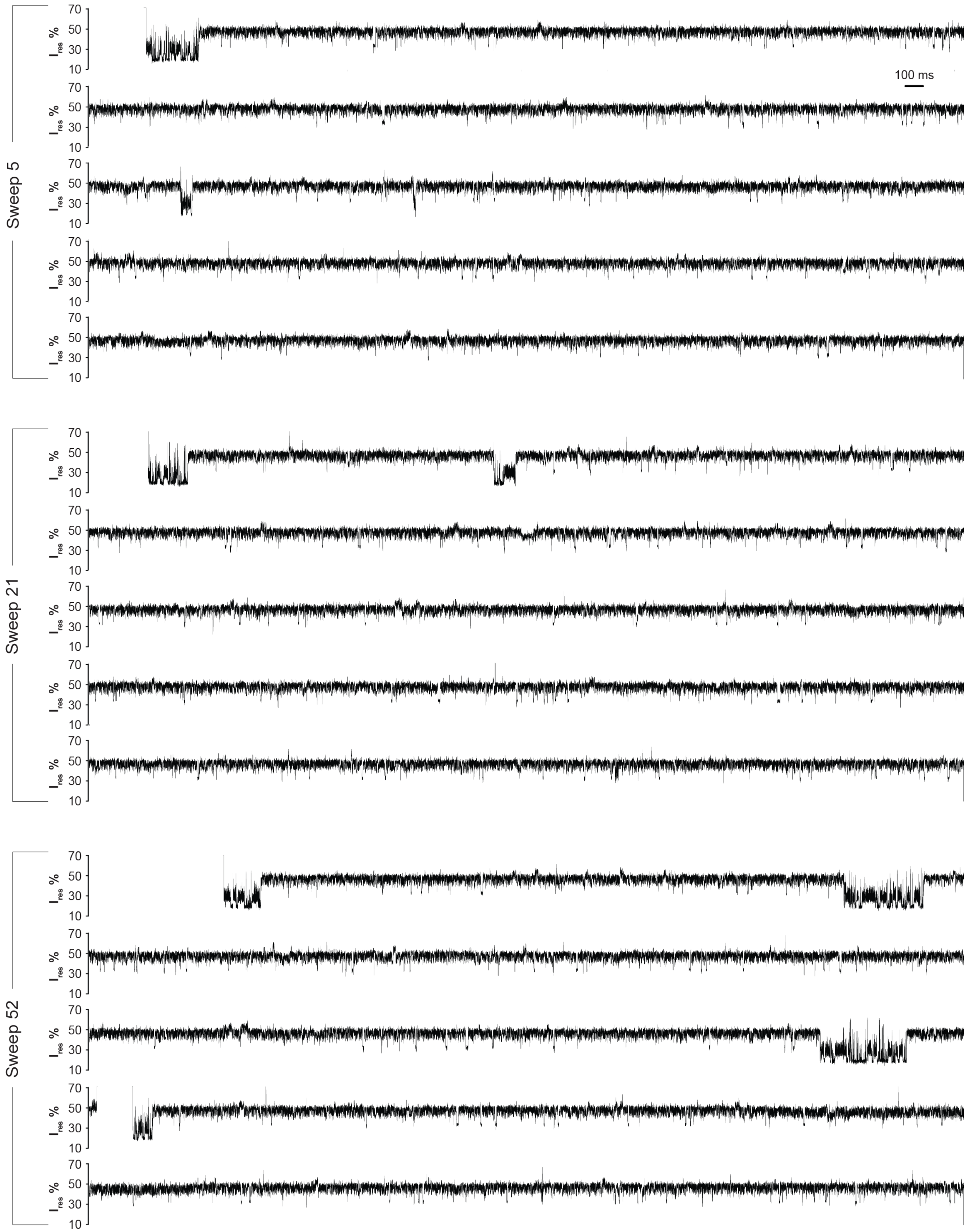


**Figure S11.** Three representative current traces Apo Abl E255V recorded using the sweep protocol. Each trace spans 25 s and was obtained at a trapping voltage of −80 mV. The $I_{res}$% range (10–70%) is shown to highlight S1/S2 signal patterns. Open-pore current and the zero baseline fall outside the displayed $I_{res}$ range and therefore appear as blank regions. Recordings were performed in 100 mM Tris-HCl (pH 7.5), 150 mM NaCl, 10 mM $MgCl_2$, and 1 mM DTT.

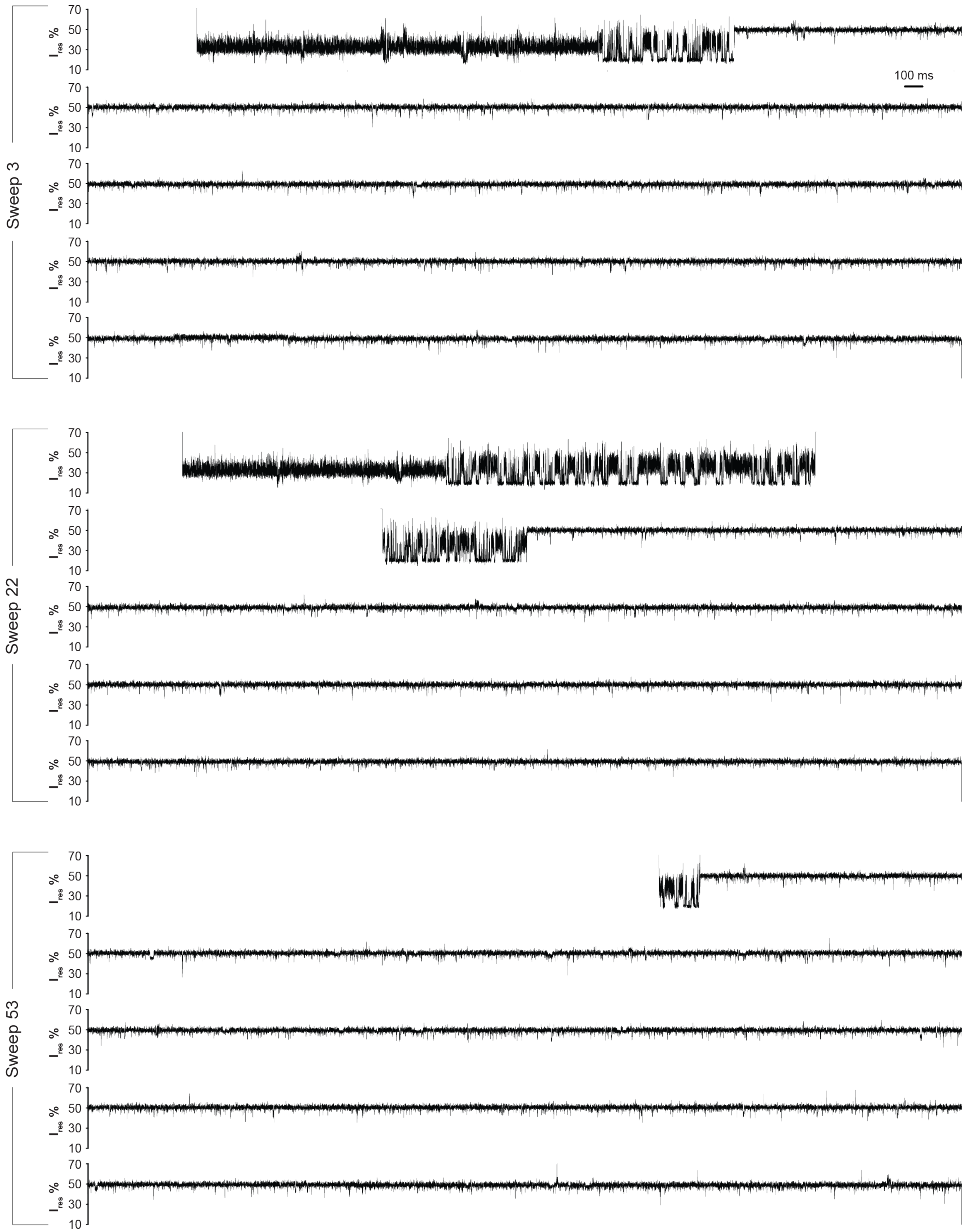


**Figure S12.** Three representative current traces of dasatinib-dosed Abl T315I recorded using the sweep protocol. Each trace spans 25 s and was obtained at a trapping voltage of −80 mV. The $I_{res}$% range (10–70%) is shown to highlight S1/S2 signal patterns. Open-pore current and the zero baseline fall outside the displayed $I_{res}$ range and therefore appear as blank regions. Recordings were performed in 100 mM Tris-HCl (pH 7.5), 150 mM NaCl, 10 mM $MgCl_2$, and 1 mM DTT, supplemented with 150 nM dasatinib at 0.15% (v/v) DMSO (final concentrations).

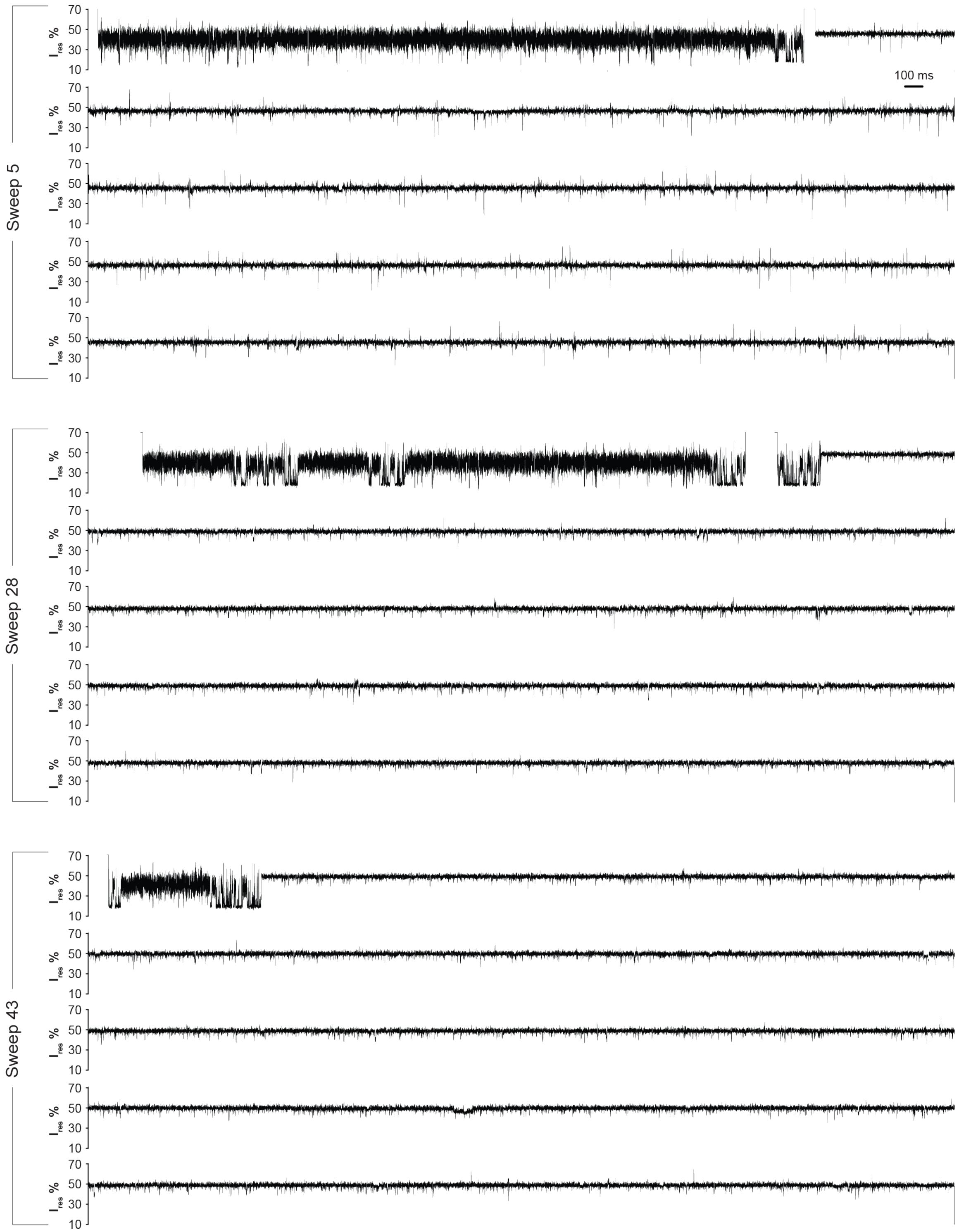


**Figure S13.** Three representative current traces of vandatenib-dosed Abl T315I recorded using the sweep protocol. Each trace spans 25 s and was obtained at a trapping voltage of −80 mV. The $I_{res}$% range (10–70%) is shown to highlight S1/S2 signal patterns. Open-pore current and the zero baseline fall outside the displayed $I_{res}$ range and therefore appear as blank regions. Recordings were performed in 100 mM Tris-HCl (pH 7.5), 150 mM NaCl, 10 mM $MgCl_2$, and 1 mM DTT, supplemented with 150 nM vandatenib at 0.15% (v/v) DMSO (final concentrations).

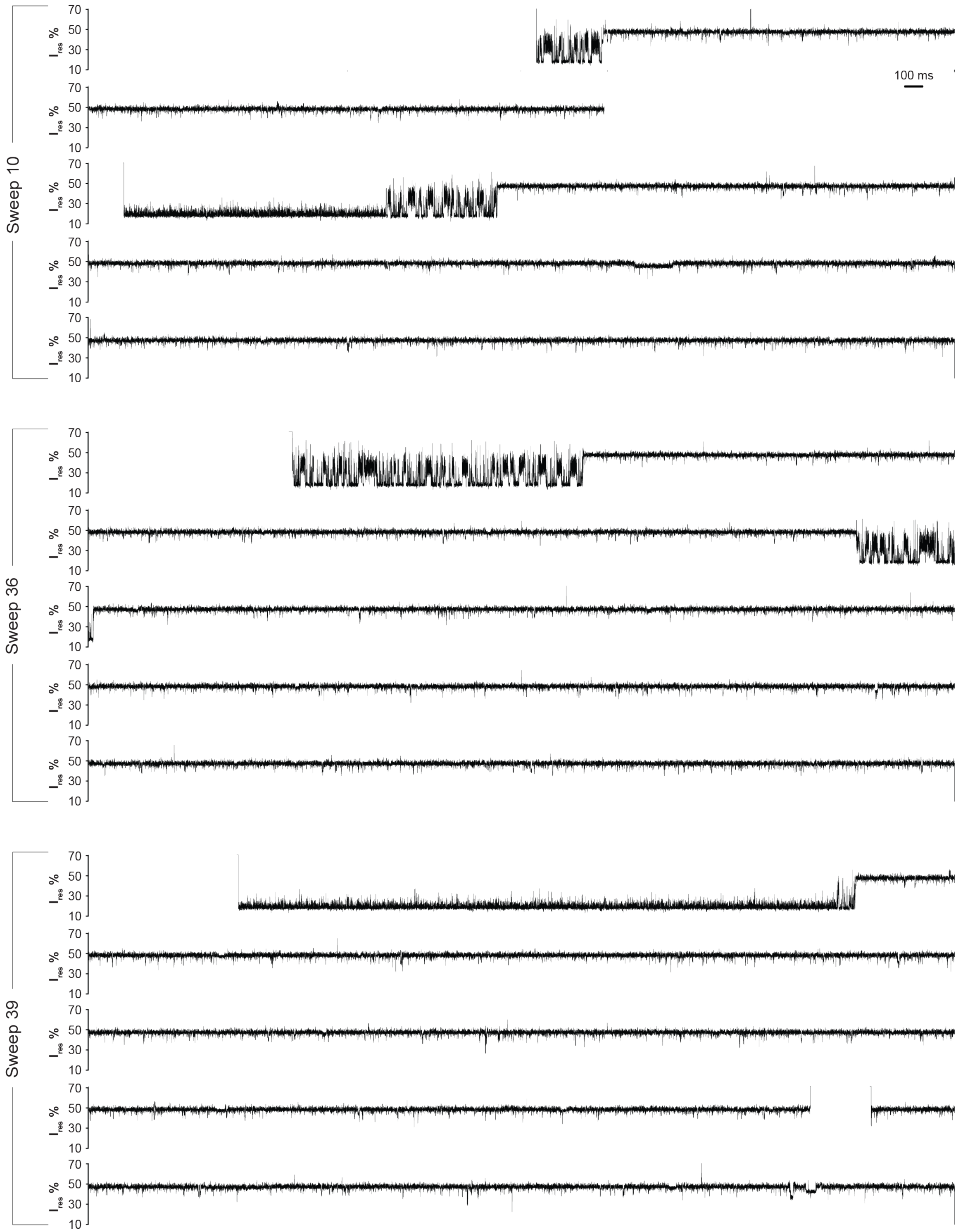


**Figure S14.** Three representative current traces of imatinib-dosed Abl T315I recorded using the sweep protocol. Each trace spans 25 s and was obtained at a trapping voltage of −80 mV. The $I_{res}$% range (10–70%) is shown to highlight S1/S2 signal patterns. Open-pore current and the zero baseline fall outside the displayed $I_{res}$ range and therefore appear as blank regions. Recordings were performed in 100 mM Tris-HCl (pH 7.5), 150 mM NaCl, 10 mM $MgCl_2$, and 1 mM DTT, supplemented with 150 nM imatinib at 0.15% (v/v) DMSO (final concentrations).

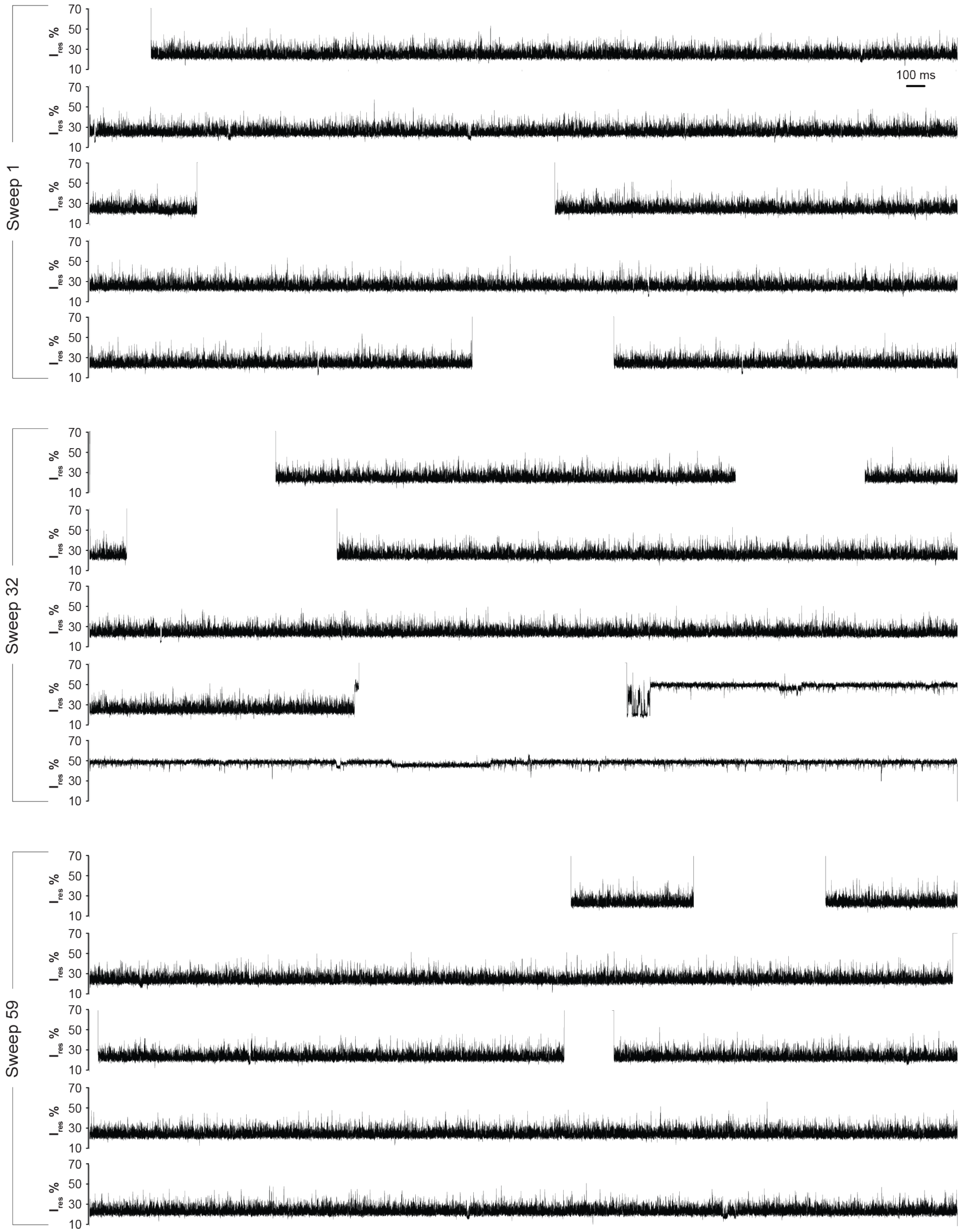


**Figure S15.** Three representative current traces of nilotinib-dosed Abl T315I recorded using the sweep protocol. Each trace spans 25 s and was obtained at a trapping voltage of −80 mV. The $I_{res}$% range (10–70%) is shown to highlight S1/S2 signal patterns. Open-pore current and the zero baseline fall outside the displayed $I_{res}$ range and therefore appear as blank regions. Recordings were performed in 100 mM Tris-HCl (pH 7.5), 150 mM NaCl, 10 mM $MgCl_2$, and 1 mM DTT, supplemented with 150 nM nilotinib at 0.15% (v/v) DMSO (final concentrations).

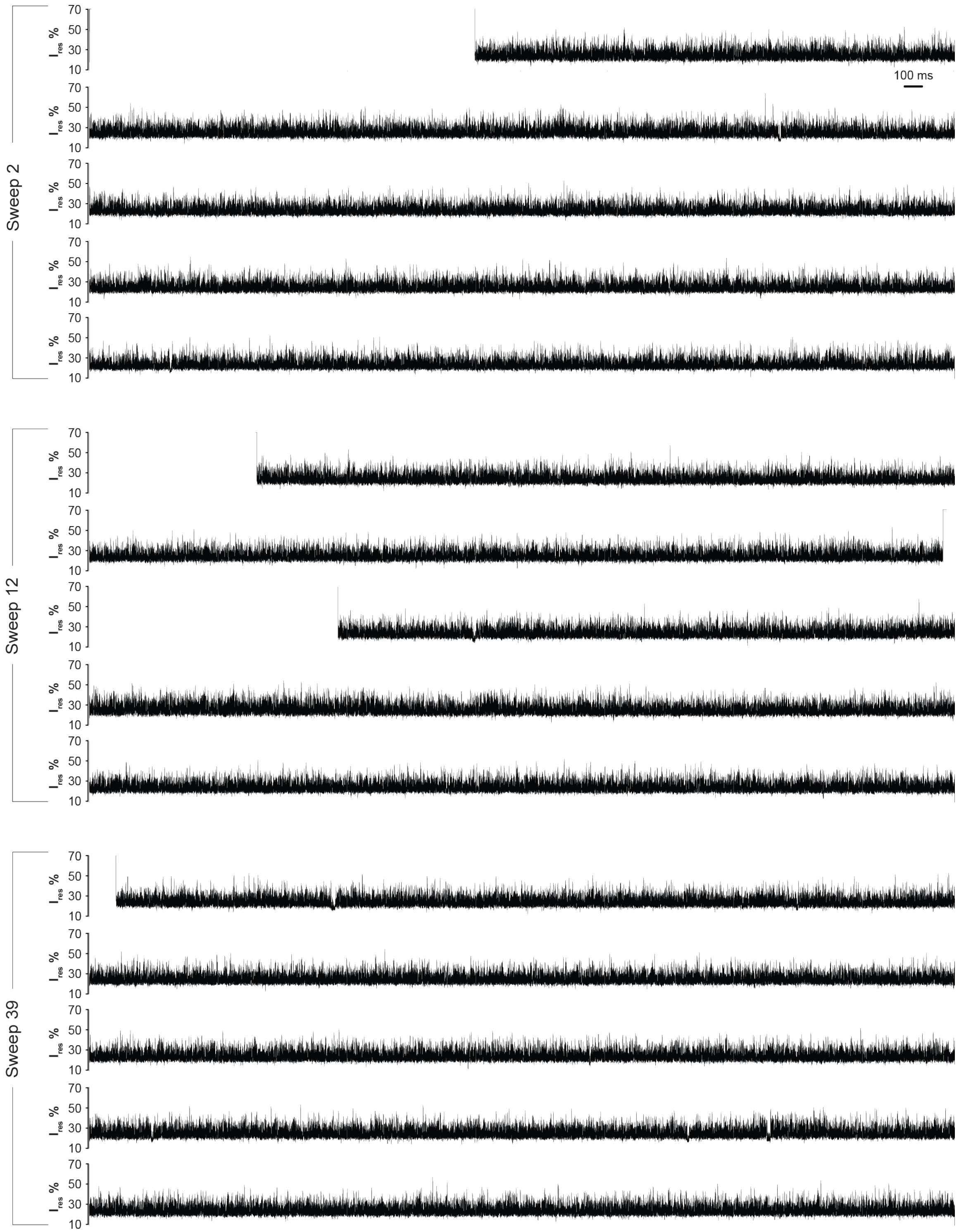


**Figure S16.** Three representative current traces of ponatinib-dosed Abl T315I recorded using the sweep protocol. Each trace spans 25 s and was obtained at a trapping voltage of −80 mV. The $I_{res}$% range (10–70%) is shown to highlight S1/S2 signal patterns. Open-pore current and the zero baseline fall outside the displayed $I_{res}$ range and therefore appear as blank regions. Recordings were performed in 100 mM Tris-HCl (pH 7.5), 150 mM NaCl, 10 mM $MgCl_2$, and 1 mM DTT, supplemented with 150 nM ponatinib at 0.15% (v/v) DMSO (final concentrations).

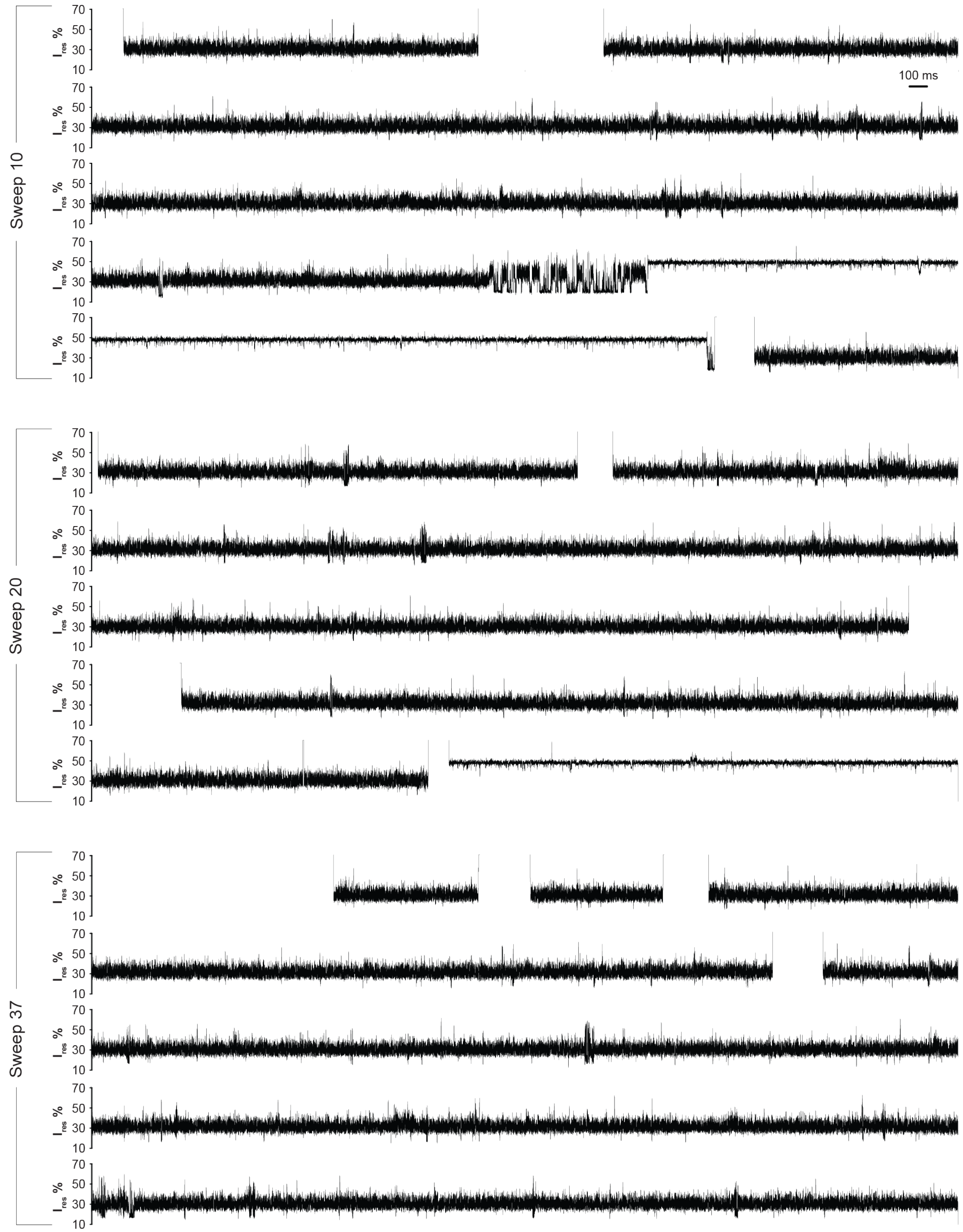


**Figure S17.** Three representative current traces of bosutinib-dosed Abl T315I recorded using the sweep protocol. Each trace spans 25 s and was obtained at a trapping voltage of −80 mV. The $I_{res}$% range (10–70%) is shown to highlight S1/S2 signal patterns. Open-pore current and the zero baseline fall outside the displayed $I_{res}$ range and therefore appear as blank regions. Recordings were performed in 100 mM Tris-HCl (pH 7.5), 150 mM NaCl, 10 mM $MgCl_2$, and 1 mM DTT, supplemented with 150 nM bosutinib at 0.15% (v/v) DMSO (final concentrations).

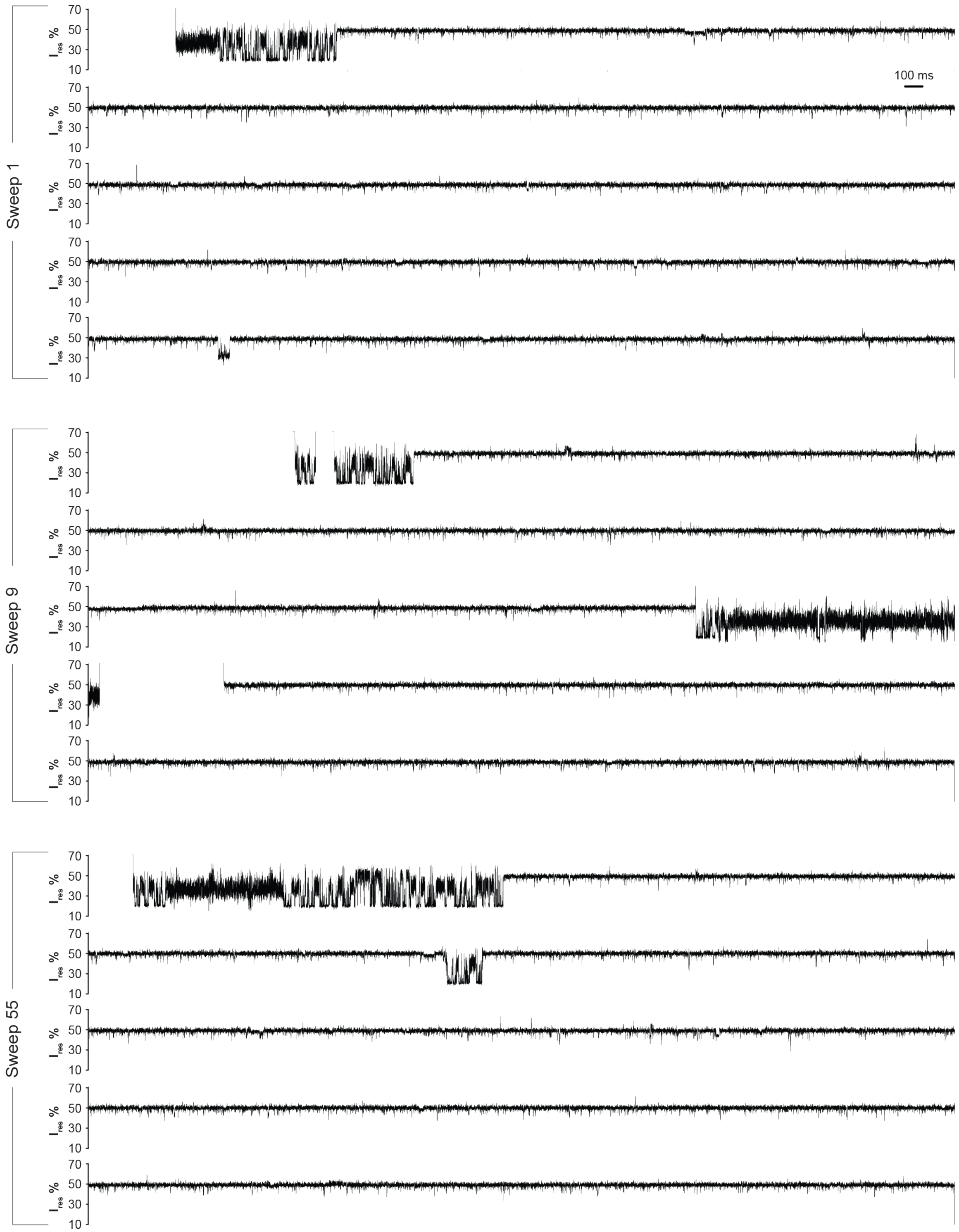


**Figure S18.** Three representative current traces of saracatinib-dosed Abl T315I recorded using the sweep protocol. Each trace spans 25 s and was obtained at a trapping voltage of −80 mV. The $I_{res}$% range (10–70%) is shown to highlight S1/S2 signal patterns. Open-pore current and the zero baseline fall outside the displayed $I_{res}$ range and therefore appear as blank regions. Recordings were performed in 100 mM Tris-HCl (pH 7.5), 150 mM NaCl, 10 mM $MgCl_2$, and 1 mM DTT, supplemented with 150 nM saracatinib at 0.15% (v/v) DMSO (final concentrations).

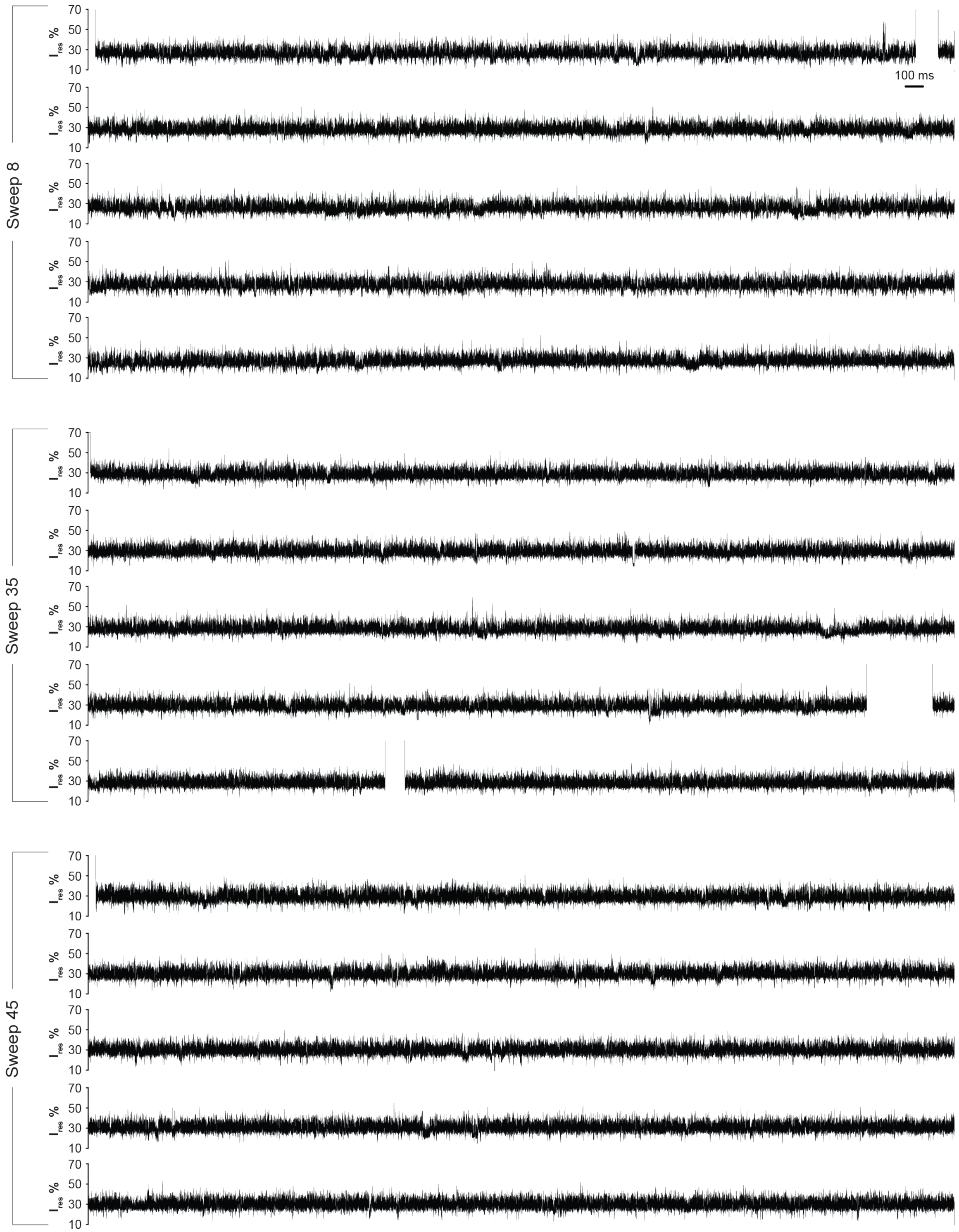


**Figure S19.** Three representative current traces of dasatinib-dosed Abl E255V recorded using the sweep protocol. Each trace spans 25 s and was obtained at a trapping voltage of −80 mV. The $I_{res}$% range (10–70%) is shown to highlight S1/S2 signal patterns. Open-pore current and the zero baseline fall outside the displayed $I_{res}$ range and therefore appear as blank regions. Recordings were performed in 100 mM Tris-HCl (pH 7.5), 150 mM NaCl, 10 mM $MgCl_2$, and 1 mM DTT, supplemented with 150 nM dasatinib at 0.15% (v/v) DMSO (final concentrations).

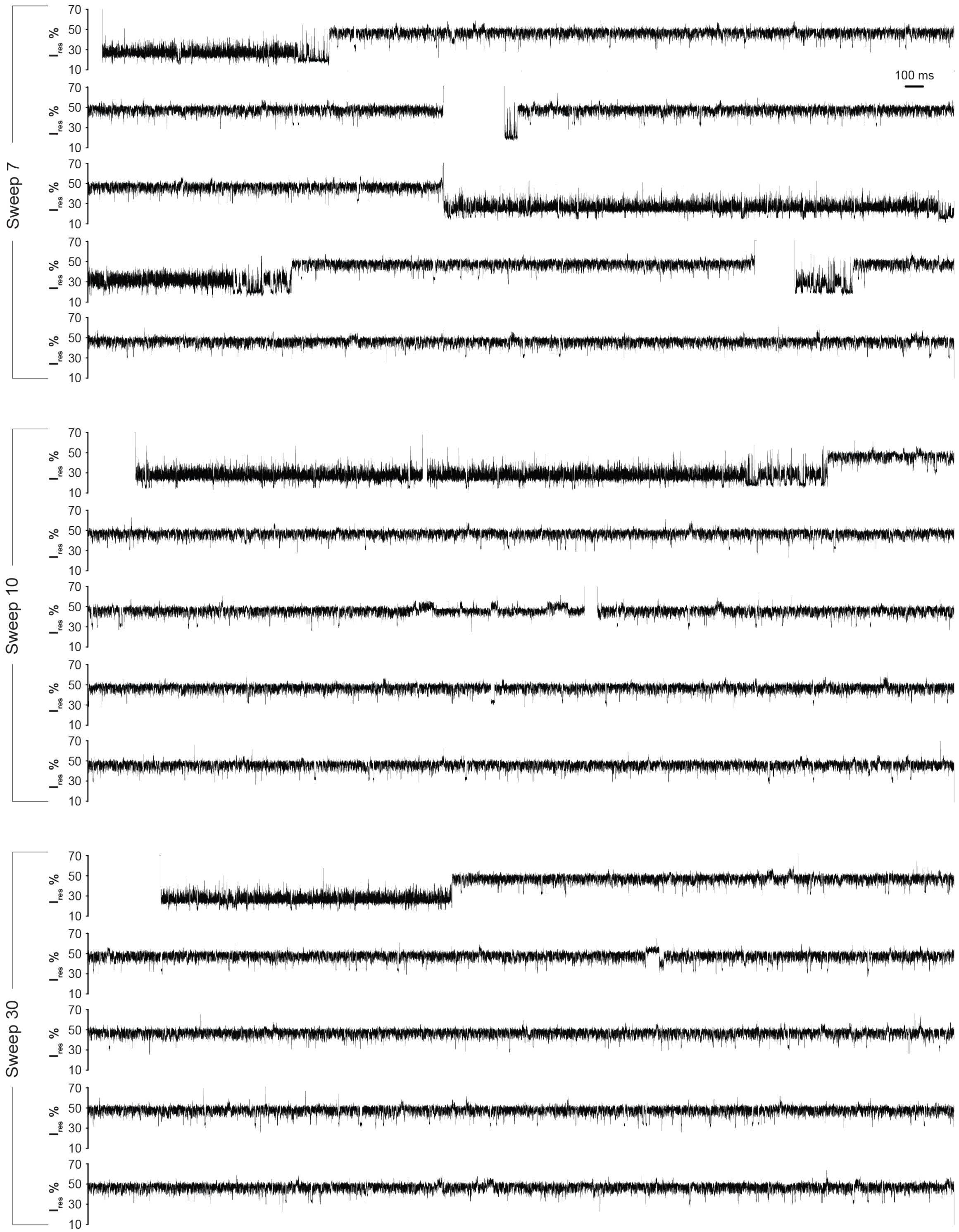


**Figure S20.** Three representative current traces of vandatenib-dosed Abl E255V recorded using the sweep protocol. Each trace spans 25 s and was obtained at a trapping voltage of −80 mV. The $I_{res}$% range (10–70%) is shown to highlight S1/S2 signal patterns. Open-pore current and the zero baseline fall outside the displayed $I_{res}$ range and therefore appear as blank regions. Recordings were performed in 100 mM Tris-HCl (pH 7.5), 150 mM NaCl, 10 mM $MgCl_2$, and 1 mM DTT, supplemented with 150 nM vandatenib at 0.15% (v/v) DMSO (final concentrations).

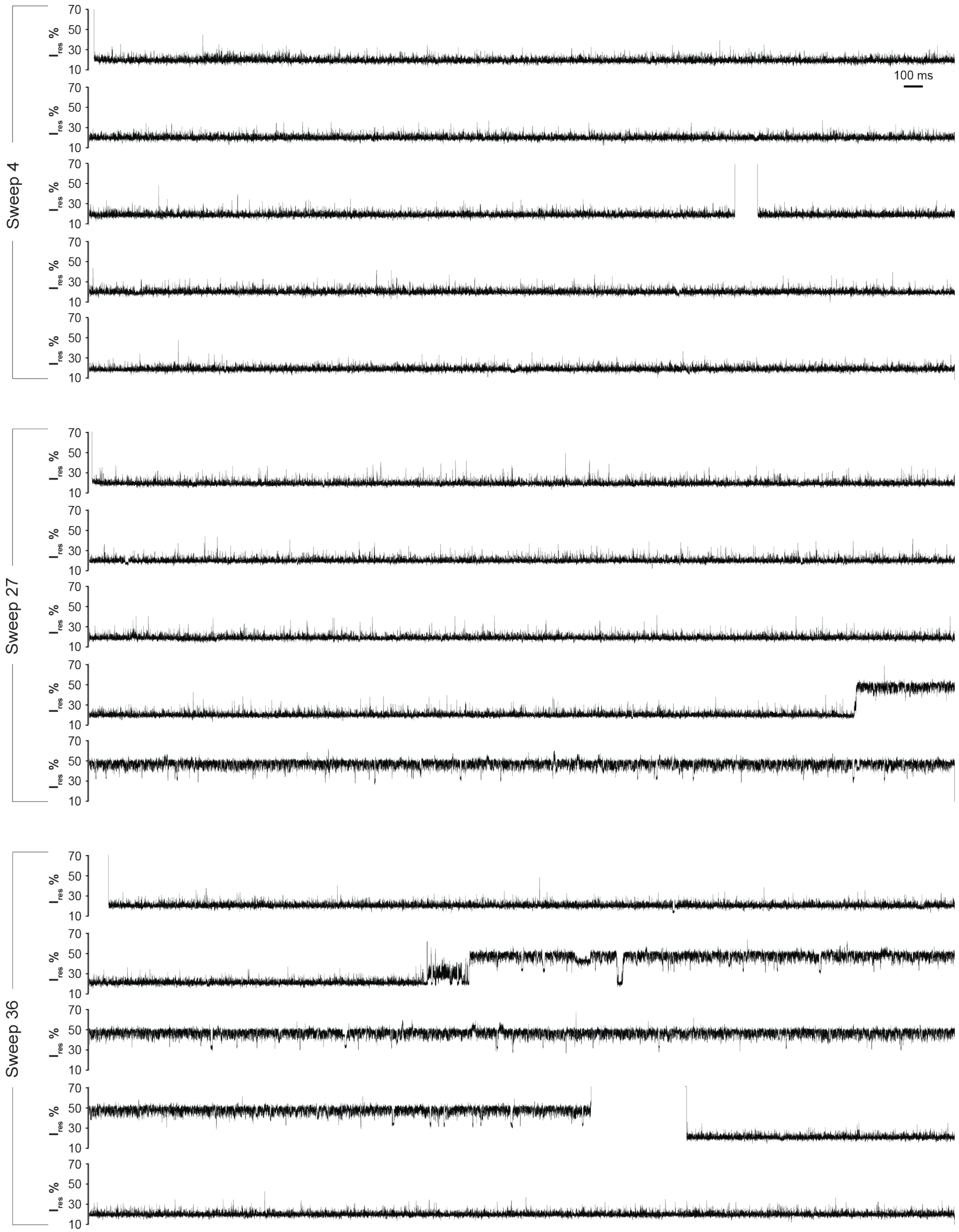


**Figure S21.** Three representative current traces of imatinib-dosed Abl E255V recorded using the sweep protocol. Each trace spans 25 s and was obtained at a trapping voltage of −80 mV. The $I_{res}$% range (10–70%) is shown to highlight S1/S2 signal patterns. Open-pore current and the zero baseline fall outside the displayed $I_{res}$ range and therefore appear as blank regions. Recordings were performed in 100 mM Tris-HCl (pH 7.5), 150 mM NaCl, 10 mM $MgCl_2$, and 1 mM DTT, supplemented with 150 nM imatinib at 0.15% (v/v) DMSO (final concentrations).

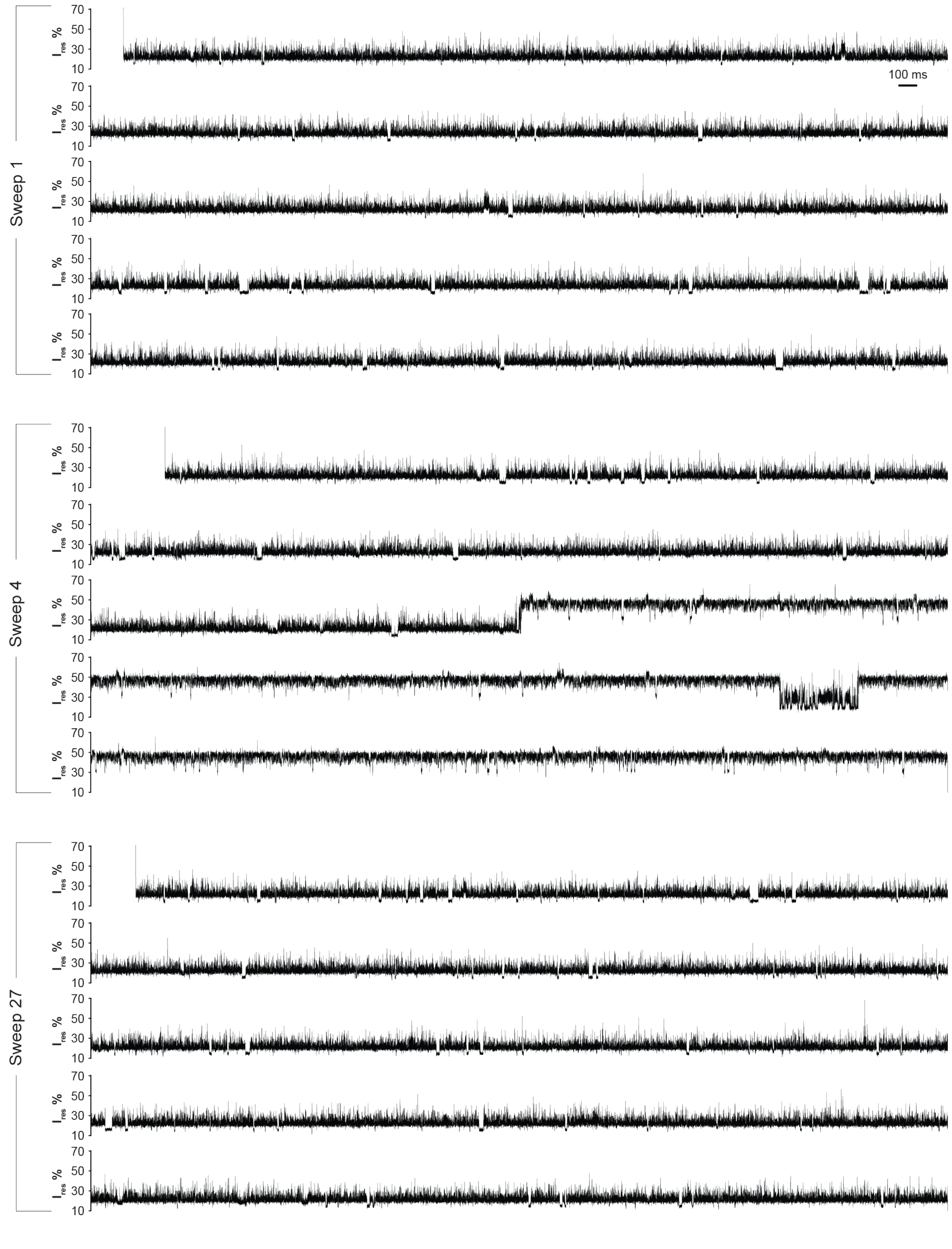


**Figure S22.** Three representative current traces of nilotinib-dosed Abl E255V recorded using the sweep protocol. Each trace spans 25 s and was obtained at a trapping voltage of −80 mV. The $I_{res}$% range (10–70%) is shown to highlight S1/S2 signal patterns. Open-pore current and the zero baseline fall outside the displayed $I_{res}$ range and therefore appear as blank regions. Recordings were performed in 100 mM Tris-HCl (pH 7.5), 150 mM NaCl, 10 mM $MgCl_2$, and 1 mM DTT, supplemented with 150 nM nilotinib at 0.15% (v/v) DMSO (final concentrations).

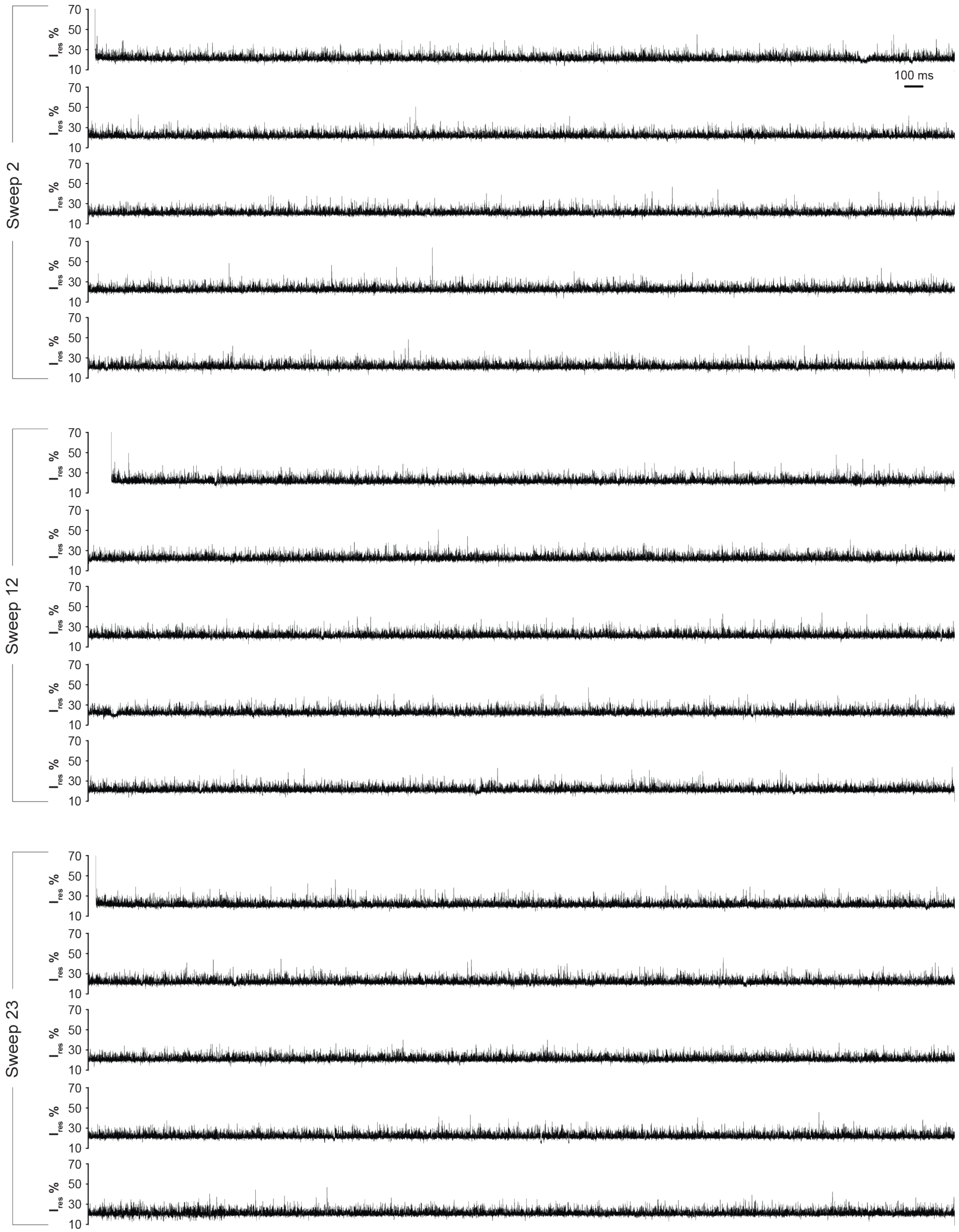


**Figure S23.** Three representative current traces of ponatinib-dosed Abl E255V recorded using the sweep protocol. Each trace spans 25 s and was obtained at a trapping voltage of −80 mV. The $I_{res}$% range (10–70%) is shown to highlight S1/S2 signal patterns. Open-pore current and the zero baseline fall outside the displayed $I_{res}$ range and therefore appear as blank regions. Recordings were performed in 100 mM Tris-HCl (pH 7.5), 150 mM NaCl, 10 mM $MgCl_2$, and 1 mM DTT, supplemented with 150 nM ponatinib at 0.15% (v/v) DMSO (final concentrations).

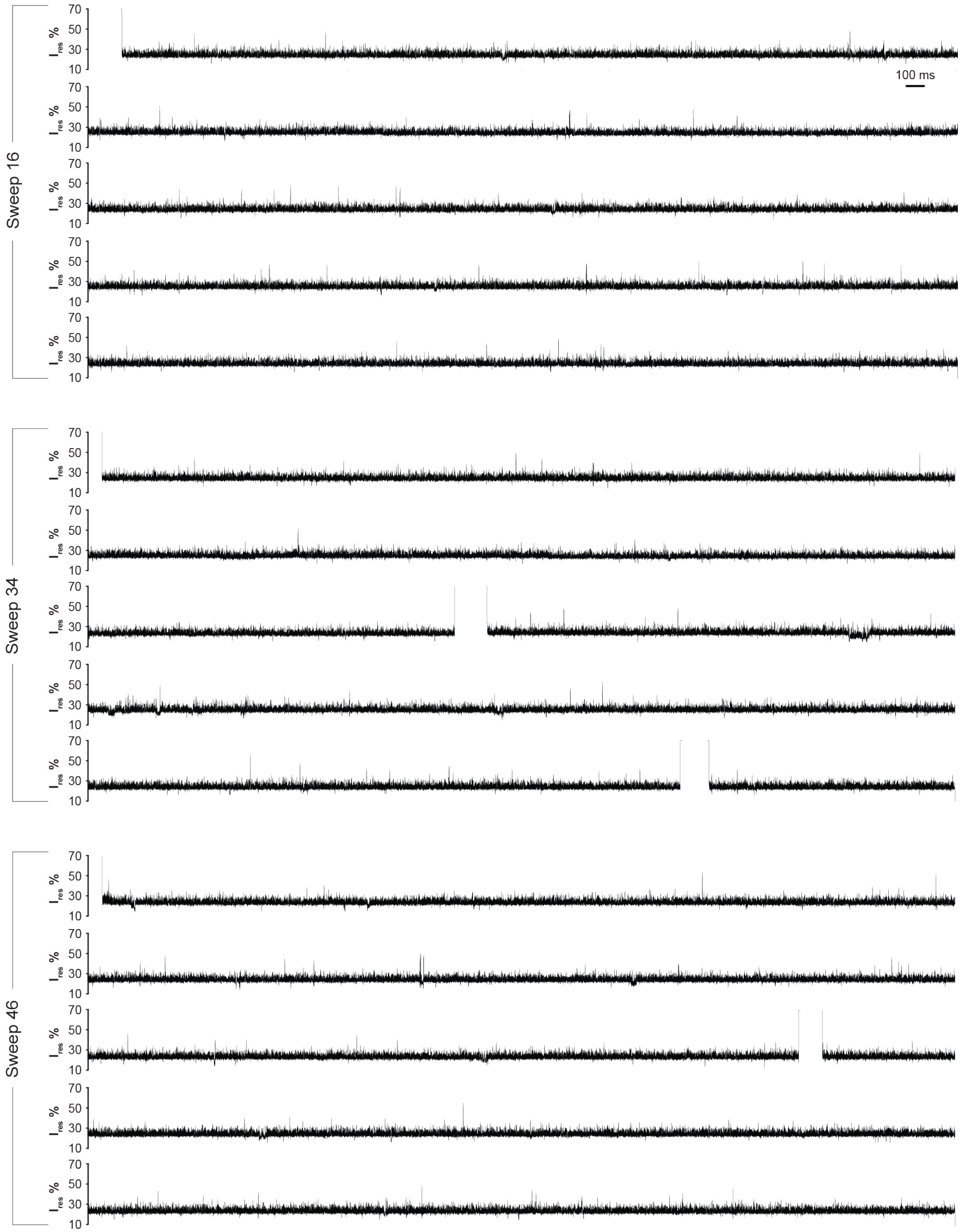


**Figure S24.** Three representative current traces of bosutinib-dosed Abl E255V recorded using the sweep protocol. Each trace spans 25 s and was obtained at a trapping voltage of −80 mV. The $I_{res}$% range (10–70%) is shown to highlight S1/S2 signal patterns. Open-pore current and the zero baseline fall outside the displayed $I_{res}$ range and therefore appear as blank regions. Recordings were performed in 100 mM Tris-HCl (pH 7.5), 150 mM NaCl, 10 mM $MgCl_2$, and 1 mM DTT, supplemented with 150 nM bosutinib at 0.15% (v/v) DMSO (final concentrations).

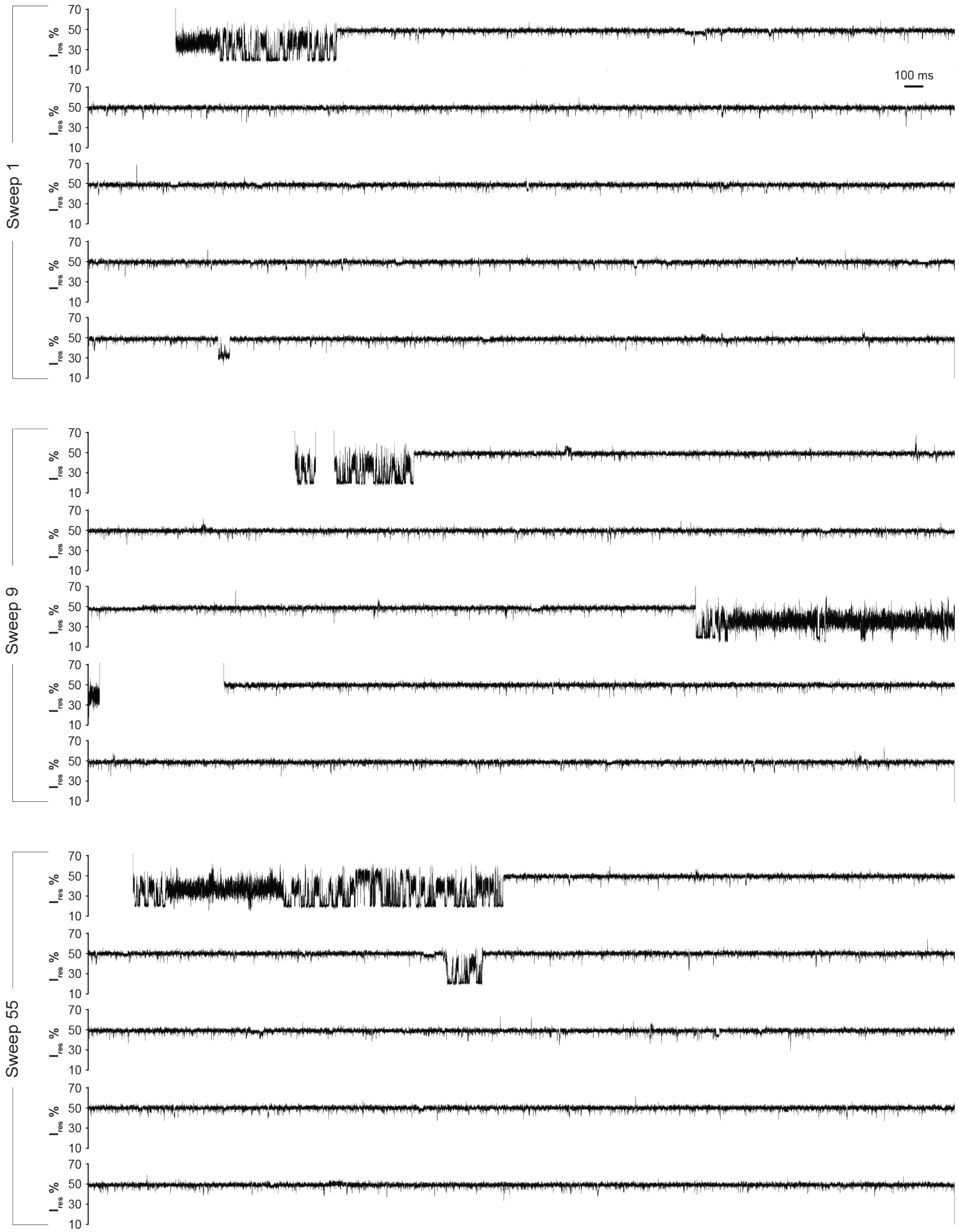


**Figure S25.** Three representative current traces of saracatinib-dosed Abl E255V recorded using the sweep protocol. Each trace spans 25 s and was obtained at a trapping voltage of −80 mV. The $I_{res}$% range (10–70%) is shown to highlight S1/S2 signal patterns. Open-pore current and the zero baseline fall outside the displayed $I_{res}$ range and therefore appear as blank regions. Recordings were performed in 100 mM Tris-HCl (pH 7.5), 150 mM NaCl, 10 mM $MgCl_2$, and 1 mM DTT, supplemented with 150 nM saracatinib at 0.15% (v/v) DMSO (final concentrations).

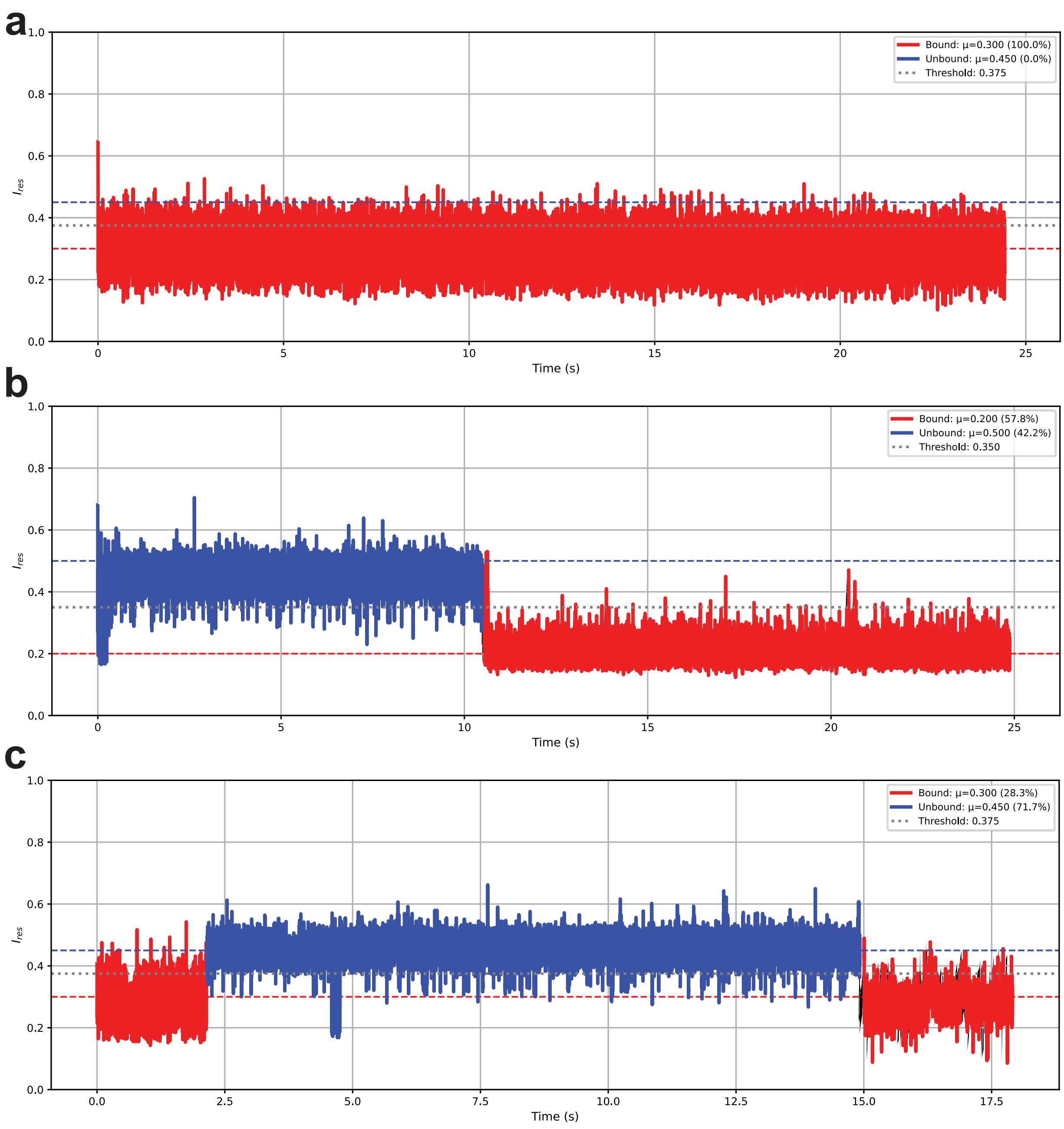


**Figure S26. Example inhibitor bound/unbound analysis traces.** Example assignments of inhibitor bound and unbound portions for capture events of Abl E255V in solution with dasatinib (a), imatinib (b), and vanatinib (c) are shown. For each trace, the inhibitor-bound signal is shaded in red, and the inhibitor-unbound signal is shaded in blue. The dashed red and blue lines represent the empirically determined Ires references used for the bound and unbound signals respectively while the threshold midpoint between the bound/unbound references is represented by the dotted grey line. The percentage of the capture event belonging to inhibitor-bound/unbound is listed in the subfigure's legends.

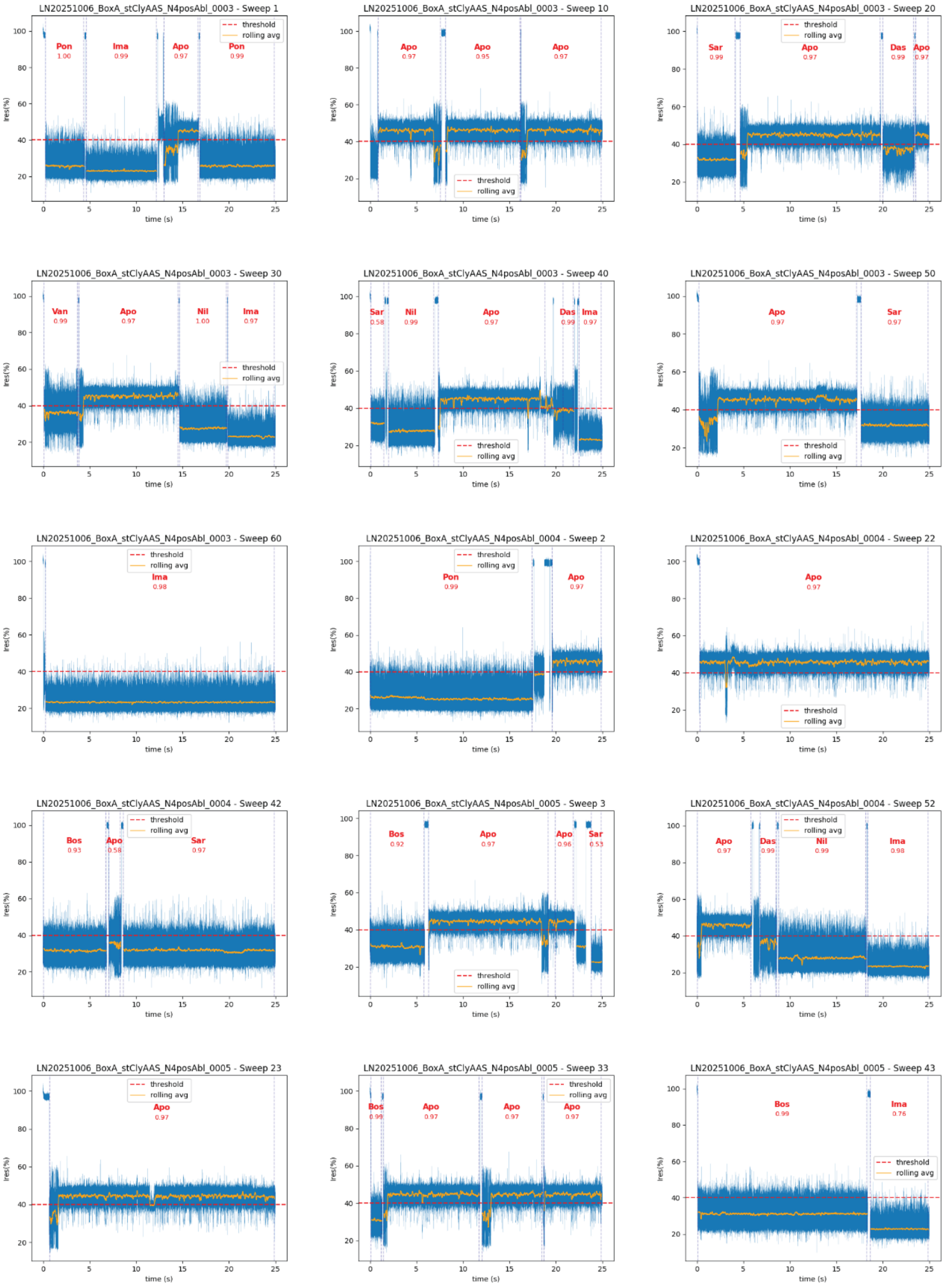


**Figure S27.** Fifteen representative sweeps from Mixture A, which contains all seven studied inhibitors (6 nM each) dosed to Abl WT, are shown along with the corresponding predicted results and confidence scores. The mixture yielded 465 individual events from 180 sweeps, including both apo and inhibitor-bound segments.

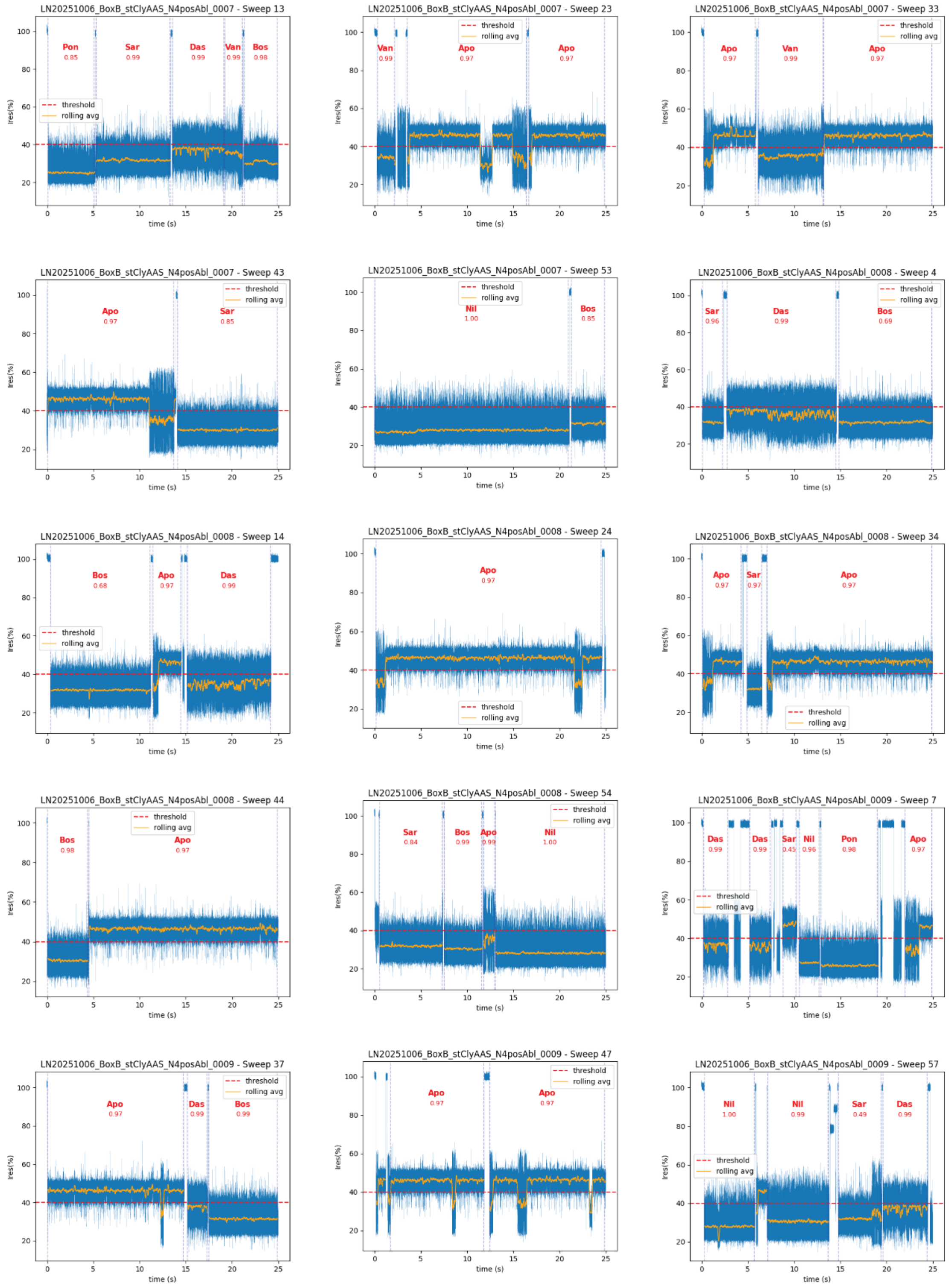


**Figure S28.** Fifteen representative sweeps from Mixture B1, which contains six studied inhibitors (6 nM each) excluding imatinib dosed to Abl WT, are shown along with the corresponding predicted results and confidence scores. The mixture yielded 474 individual events from 180 sweeps, including both apo and inhibitor-bound segments.

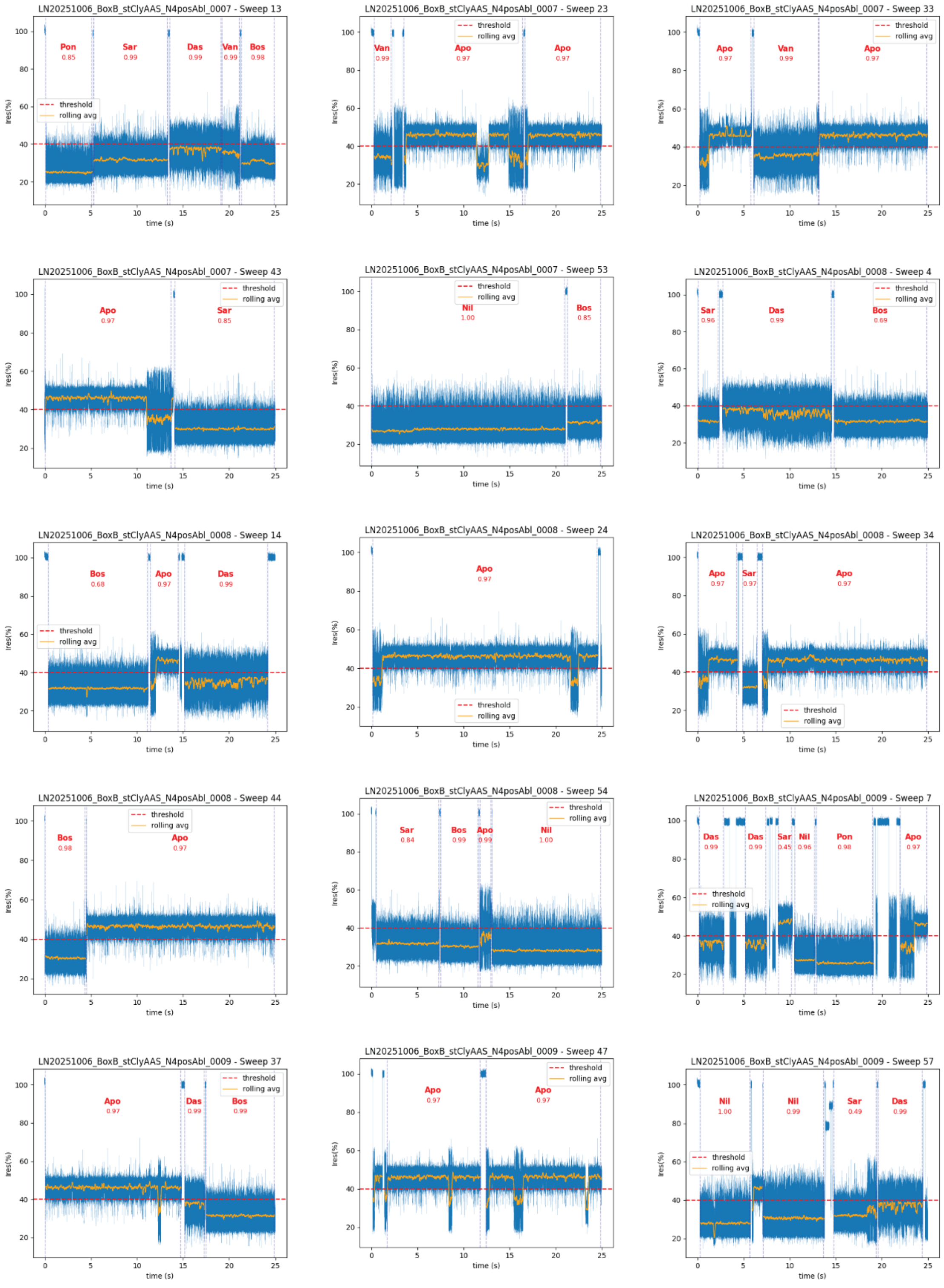


**Figure S29.** Fifteen representative sweeps from Mixture B2, which contains six studied inhibitors (6 nM each) excluding bosutinib dosed to Abl WT, are shown along with the corresponding predicted results and confidence scores. The mixture yielded 464 individual events from 180 sweeps, including both apo and inhibitor-bound segments.

**Table S1.** Quantitative analysis results for Inhibitor Mixtures Dosed to Abl WT

| | **Mixture A** | | | | |
|---|---|---|---|---|---|
| **Inhibitor** | Count | Sum Length (s) | **% Bound** | Avg Length (s) | Std Length |
| Dasatinib | 44 | 156.65 | 3.74 | 3.56 | 2.64 |
| Vandetanib | 21 | 56.54 | 1.35 | 2.69 | 1.11 |
| Imatinib | 47 | 674.53 | 16.11 | 14.35 | 8.56 |
| Nilotinib | 37 | 357.83 | 8.55 | 9.67 | 6.54 |
| Ponatinib | 38 | 370.43 | 8.85 | 9.75 | 7.45 |
| Bosutinib | 65 | 535.93 | 12.80 | 8.25 | 6.01 |
| Saracatinib | 45 | 305.37 | 7.29 | 6.79 | 5.48 |
| Apo | 168 | 1728.88 | 41.30 | 10.29 | 7.36 |
| **Total** | **465** | **4186.16** | | | |

| | **Mixture B1 (- Imatinib)** | | | | |
|---|---|---|---|---|---|
| **Inhibitor** | Count | Sum Length (s) | **% Bound** | Avg Length(s) | Std Length |
| Dasatinib | 60 | 292.32 | 7.12 | 4.87 | 3.98 |
| Vandetanib | 18 | 58.43 | 1.42 | 3.25 | 1.94 |
| Imatinib | 2 | 13.90 | 0.34 | 6.95 | 5.83 |
| Nilotinib | 49 | 503.27 | 12.27 | 10.27 | 6.72 |
| Ponatinib | 44 | 452.72 | 11.03 | 10.29 | 7.35 |
| Bosutinib | 67 | 565.32 | 13.78 | 8.44 | 6.91 |
| Saracatinib | 78 | 567.89 | 13.84 | 7.28 | 6.01 |
| Apo | 156 | 1648.92 | 40.19 | 10.57 | 7.76 |
| **Total** | **474** | **4102.77** | | | |

| | **Mixture B2 (- Bosutinib)** | | | | |
|---|---|---|---|---|---|
| **Inhibitor** | Count | Sum Length (s) | **% Bound** | Avg Length(s) | Std Length |
| Dasatinib | 46 | 188.68 | 4.54 | 4.1 | 2.9 |
| Vandetanib | 20 | 63.81 | 1.53 | 3.19 | 1.82 |
| Imatinib | 57 | 791.87 | 19.04 | 13.89 | 8.46 |
| Nilotinib | 44 | 403.59 | 9.70 | 9.17 | 7.07 |
| Ponatinib | 45 | 382.37 | 9.19 | 8.5 | 6.51 |
| Bosutinib | 7 | 46.20 | 1.11 | 6.6 | 6.14 |
| Saracatinib | 54 | 325.88 | 7.84 | 6.03 | 5.3 |
| Apo | 191 | 1956.62 | 47.05 | 10.24 | 7.24 |
| **Total** | **464** | **4159.02** | | | |

**Table S2.** Primers used for generating Abl variants by site-directed mutagenesis.

| Primer | Sequence (5′→3′) | Purpose |
|---|---|---|
| T315I–F | ATCATCATTGAGTTCATGACCTACGGGAAC | Introduce the T315I mutation into the N4posAbl construct |
| T315I–R | GAACTCAATGATGATATAGAACGGGGGCTC | Introduce the T315I mutation into the N4posAbl construct |
| E255V–F | GGCCAGTACGGGGTAGTGTACGAGGGC | Introduce the E255V mutation into the N4posAbl construct |
| E255V–R | TACCCCGTACTGGCCCCCGCCCAG | Introduce the E255V mutation into the N4posAbl construct |

### References

1. Xie, T., Saleh, T., Rossi, P. & Kalodimos, C. G. Conformational states dynamically populated by a kinase determine its function. *Science* **370**, eabc2754 (2020).
